# Altered AP-1, RUNX and EGR chromatin dynamics drive fibrotic lung disease

**DOI:** 10.1101/2024.10.23.619858

**Authors:** Eleanor Valenzi, Minxue Jia, Peter Gerges, Jingyu Fan, Tracy Tabib, Rithika Behara, Yuechen Zhou, John Sembrat, Jishnu Das, Panayiotis V Benos, Harinder Singh, Robert Lafyatis

## Abstract

Pulmonary fibrosis, including systemic sclerosis-associated interstitial lung disease (SSc-ILD), involves myofibroblasts and SPP1^hi^ macrophages as drivers of fibrosis. Single-cell RNA sequencing has delineated fibroblast and macrophages transcriptomes, but limited insight into transcriptional control of profibrotic gene programs. To address this challenge, we analyzed multiomic snATAC/snRNA-seq on explanted SSc-ILD and donor control lungs. The neural network tool ChromBPNet inferred increased TF binding at single base pair resolution to profibrotic genes, including CTHRC1 and ADAM12, in fibroblasts and SPP1 and CCL18 in macrophages. The novel algorithm HALO confirmed AP-1, RUNX, and EGR TF activity controlling profibrotic gene programs and established TF-regulatory element-gene networks. This TF action atlas provides comprehensive insights into the transcriptional regulation of fibroblasts and macrophages in healthy and fibrotic human lungs.

## Introduction

Fibrosis is a lethal feature in numerous diseases. Myofibroblasts drive fibrosis due to their excessive production of extracellular matrix, resulting in tissue stiffness and organ dysfunction. While necessary for normal physiologic wound healing, dysregulated myofibroblasts contribute to fibrosis in pulmonary, renal, dermal, hepatic, and cardiac tissues. Macrophages, particularly a subset marked by SPP1, are implicated in activating myofibroblasts and infiltrate fibrotic tissues, including the lungs, liver^1^, skeletal muscle^2^ and heart^3–5^. Deleting or silencing *SPP1*, encoding Osteopontin, ameliorates bleomycin induced lung fibrosis^6–8^. Single cell RNA-sequencing (scRNA-seq) meta-analyses reveal that SPP1^hi^ macrophages share similar transcriptional states across organs affected by fibrosis^1,9^.

Systemic sclerosis is a prototypic fibrotic disease, with fibrosis affecting multiple organs. Systemic sclerosis-associated interstitial lung disease (SSc-ILD) is the leading cause of disease-associated mortality in patients with SSc^10^. Despite current therapies targeting immune dysregulation, vasculopathy, and fibrosis, limited progress has been made in ameliorating disease-associated mortality or improving quality of life. Myofibroblasts and SPP1^hi^ macrophages are prominent cell populations and are associated with progressive lung disease^11^. Thus, a deeper understanding of the transcriptional programs regulating these cells’ profibrotic phenotypes is critical for elucidating the molecular mechanisms underlying disease and informing new therapeutic strategies.

ScRNA-seq has delineated the transcriptomes of various cell types in the healthy and fibrotic lung^12–14^, revealing perturbed cell states and altered cell-cell communication networks in disease^15,16,17^. Despite generating biologically relevant inferences of altered cell phenotypes and signaling pathways, these studies are limited in their ability to decipher how gene regulatory networks are perturbed, leading to the altered cellular states present in fibrosis. Gene expression is regulated in a cell type- and tissue-specific manner by transcription factors (TFs) interacting with *cis*-regulatory elements (CREs) in the genome. Understanding changes in TF dynamics at CREs in key profibrotic cell populations, namely myofibroblasts and SPP1^hi^ macrophages, can provide direct insights into disrupted regulatory networks in fibrosis.

Single-nucleus ATAC-seq (snATAC-seq) permits the identification of open (accessible) chromatin regions (OCRs), that represent candidate CREs, in diverse cell populations including from diseased tissues. ChromBPNet, a convolutional neural network model, predicts at single-base-pair resolution how TF action contributes to chromatin accessibility^18^. Perturbations in ChromBPNet contribution scores of TF motifs infer changes in TF dynamics in the context of disease. Using multiomic snATAC-seq/snRNA-seq on cells isolated from human SSc-ILD and healthy lungs, we identified differentially accessible chromatin regions and TF binding dynamics, allowing us to infer gene-regulatory networks in fibrotic SSc-ILD cell populations. Focusing on fibroblasts and SPP1^hi^ macrophages, we identified altered binding of AP-1, RUNX and EGR TFs in SSc-ILD fibroblasts, as well as basic helix-loop-helix(bHLH) ZIP and AP-1 TFs in SSc-ILD macrophages involving CREs associated with genes promoting fibrosis. These studies provide *in vivo* insights into the altered TF dynamics driving the profibrotic states of these cell populations.

## Results

### Epigenomic markers of healthy and SSc-ILD lung cell populations

We performed paired scRNA-seq and snATAC-seq on 3 SSc-ILD and 2 control lung samples, and multiome snRNA-seq/snATAC-seq on nuclei from 6 SSc-ILD and 7 control explants. Subpleural tissue was collected at lung transplant in patients with SSc-ILD, and from organ donors without preexisting lung disease. Sixteen samples were partially enriched for mesenchymal cells. Histology showed usual interstitial pneumonia in 8 SSc-ILD samples and more predominant vascular disease in 1(Supp Fig 1, Supp Table 1). After filtering, 67,446 nuclei (average 3,747 per donor) were included for analyses. Normalization, batch correction, and dimensionality reduction were performed using Seurat, Signac, and Harmony^19,20^. SnATAC data alone was utilized for initial clustering after delineating OCRs/peaks using MACS2. We identified 35 distinct clusters based on gene expression and promoter accessibility (Figure 1A-B, Supp Fig 2). All major pulmonary cell types were identified with 446,087 OCRs among 67,446 nuclei. Nuclei clustered by cell type and were well integrated by chemistry, disease state, and individual sample after batch correction (Figure 1C-D, Supp Fig 3A-B). Low quality clusters were excluded from further analyses.

**Figure 1.**
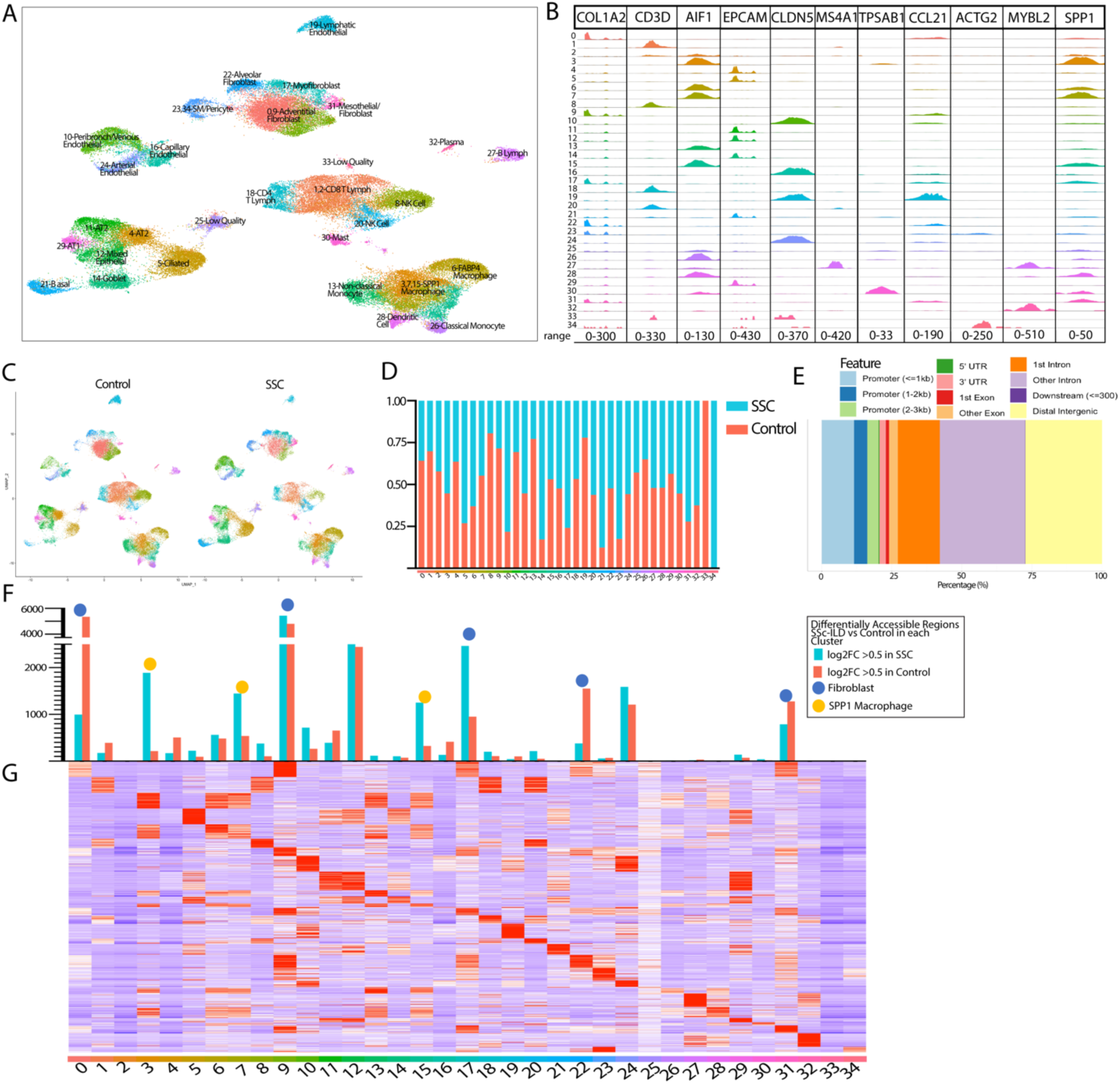
A- Clustering of the 9 SSc and 9 Control samples after filtering, includes 67, 446 nuclei. B- Identification of cell clusters by ATAC pseudobulk in promoter regions of known cell type identifying genes C- Clustering divided by Control versus SSc origin D- Proportion of cells in each cluster originating from SSc versus Control lungs E- Annotation of peak regions for lung clustering. Peaks called per cluster by macs2 with annotation by ChIPseeker F- Number of statistically significant (adj p-val <0.05) differentially accessible regions when comparing SSc versus Control nuclei within each cluster G- Top differentially accessible regions for each cluster

OCRs were annotated to the nearest gene and region (Figure 1E), with 20.32% of OCRs occurring within 3 kb of transcription start sites, and most OCRs in distal intergenic or intronic regions.^21^ Cell clusters were distinguished by differentially expressed genes (DEGs) and differentially accessible chromatin regions (DARs), with cluster 9 and 31 fibroblasts showing the most unique DARs (Figure 1G). DEGs and DARs were identified between SSc and control for each population (Figure 1F), with fibroblast, SPP1^hi^ macrophage, mixed epithelial, and arterial endothelial cells having the largest number of significant DARs, reflecting significant chromatin accessibility changes in these cell types with fibrosis.

Next, we identified cell type-specific TFs by gene expression and motif enrichment. TF motifs were identified using chromVAR^22^ (Supp Fig 4 A, B), revealing known regulators such as ETS1 in NK and T cells, SPI1 in macrophages, and GATA2 in mast cells, validating the analysis. Enriched fibroblast motifs included AP-1, Tal-related (including TWIST1 and HAND2), and TEAD TF family motifs. While TF motif activity and gene expression showed overall limited correlation, 23 TFs had significant positive correlations, including TWIST1 and HAND2 in fibroblasts, SPI1 in macrophages, and SOX18 in endothelial cells (Supp Fig 4C). We also evaluated enriched motifs for control lungs only, with similar results (Supp Fig 5).

### Altered Chromatin accessibility of fibrotic mesenchymal cell populations in SSc-ILD

Given the significant chromatin accessibility changes between SSc and control fibroblasts, we subclustered fibroblast, smooth muscle, and pericyte populations (n=13,766 nuclei; Figure 2A). We identified 10 fibroblast populations, including 4 adventitial 2 alveolar, a CXCL2^hi^ fibroblast, and 3 myofibroblast-related clusters (myofibroblast, inflammatory myofibroblast, and activated myofibroblast). Myofibroblasts, inflammatory myofibroblasts, and activated myofibroblasts were expanded in SSc-ILD, along with smooth muscle and pericytes (Figure 2B,C, Supp Fig 6A), consistent with prior scRNA-seq analyses^14^, and reflective of both severe ILD and the group 3 pulmonary hypertension present in these patients.

**Figure 2.**
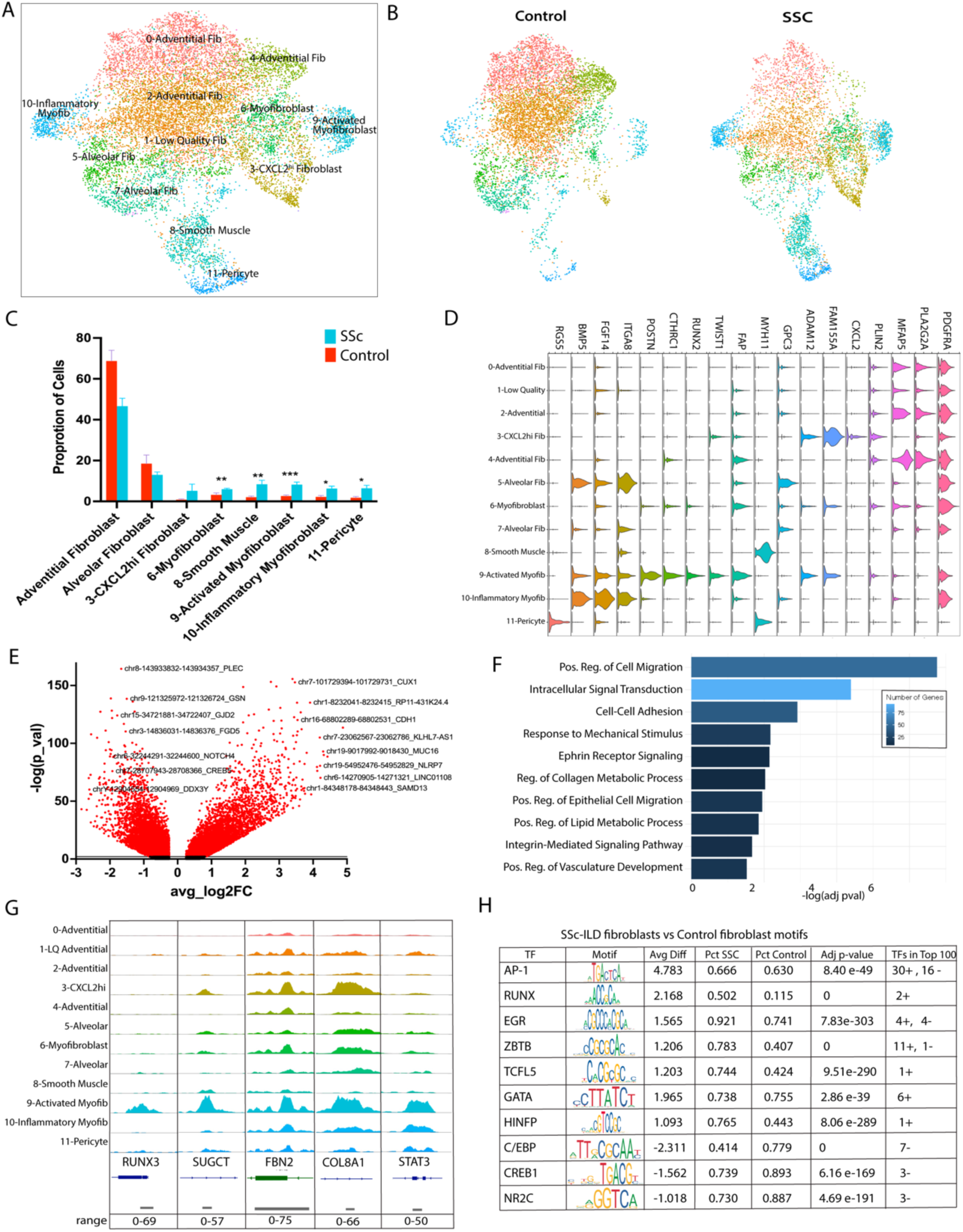
A- Subclustering of mesenchymal nuclei B- Mesenchymal nuclei divided by Control versus SSc C- Proportion of mesenchymal nuclei consisting of each cell type at the individual sample level, comparing SSc vs Control. Bar represents the mean with error bar the standard error of the mean. *=0.0106-0.0142, **=p=0.0028-0.0056, ***=p=0.0005; D- Gene expression of select cell type identifiers. E- SSc Fibroblasts vs Control Fibroblasts DARs with labeling of select DARs and the nearest annotated gene F- Pathway enrichment of OCRs more accessible in SSc vs Control fibroblasts G- ATAC pseudobulk by mesenchymal cluster of 5 regions with top DARs in activated myofibroblasts H- Top enriched TF family motifs in SSc-ILD fibroblasts vs Control fibroblasts, ranked by absolute value of average difference and separated into positively and negatively enriched. Final column notes number of TFs in the same class/family present in the top 100, as well as the direction of their enrichment (positive or negative)

We compared SSc-ILD fibroblasts to controls, SSc-ILD myofibroblasts vs SSc non-myogenic fibroblasts (including the adventitial, alveolar, and CXCL2^hi^ populations), and activated myofibroblasts to other fibroblasts to robustly investigate the DEGs in these fibroblast population (Supp Fig. 6C and 7). Activated myofibroblasts exhibited the highest expression of myofibroblast hallmark genes including *CTHRC1, POSTN, COMP,* and *COL8A1* (Fig 2D, Supp Fig 6C) and were associated with profibrotic pathways (Supp Fig 6D,E). Forty-two DEGs were upregulated in common in SSc fibroblasts, SSc myofibroblasts, and activated myofibroblasts, including *POSTN, TWIST1,* and *COL8A1* (Supp Fig 7).

In fibroblast clusters, 6,002 DARS were more accessible in SSc vs 6,693 in control fibroblasts (Figure 2E, Supp Table 2) with Cell migration and Collagen metabolic processes amongst the top upregulated pathways (Figure 2F). Activated myofibroblasts showed significant overlap in DARs with inflammatory myofibroblasts and myofibroblasts, suggesting similar epigenetic regulation (Supp Fig. 8A). Of 1,024 DARs more accessible in both SSc myofibroblasts and activated myofibroblasts (Supp Fig. 8D), 96 were annotated to upregulated genes in comparing SSc-ILD myofibroblasts vs non-myogenic fibroblasts, including *COMP*, *COL8A1*, and *TWIST.* Many shared DARs occurred in genes previously related to fibrosis such as *COL8A1, FBN2* and *STAT3* (Figure 2G).

Mapping TF motifs in OCRs of SSc vs control fibroblasts revealed that AP-1, RUNX, and EGR families had the highest positive average difference motif activity scores, while C/EBP motifs showed more accessibility in control fibroblast OCRs (Figure 2H). Since multiple related TFs often share nearly equivalent motifs, enriched TFs with adjusted p-value <0.05 and absolute average difference of 0.5 or greater were ranked based on the top score for each TF family.

### Enhanced binding of AP-1, RUNX and EGR transcription factors by fibrotic fibroblasts

To investigate chromatin accessibility dynamics in fibroblasts, we employed ChromBPNet, a deep learning-based tool trained using DNA sequences and their corresponding ATAC-seq DNA cleavage chromatin accessibility patterns, that generates base pair resolution contribution scores after removing Tn5 enzyme cutting bias^18^. After removing bias, corrected signal profiles for each cell state were used to generate single-base pair resolution contribution scores, with de novo motif discovery by TF-MoDISco^23,24^ and annotation by JASPAR TF motifs^25^. Analysis peaks were called using ChromBPNet contribution score tracks, with aligned genomic coordinates for a given TF motif used to generate cluster-specific aggregate contribution scores (Figure 3A). The seqlets inferred by ChromBPNet as showing TF binding were consistent with known binding for several families of TFs and showed differences in seqlet numbers between control and SSc-ILD fibroblasts for AP-1, RUNX, EGR, and C/EBP families (Figure 3B).

**Figure 3.**
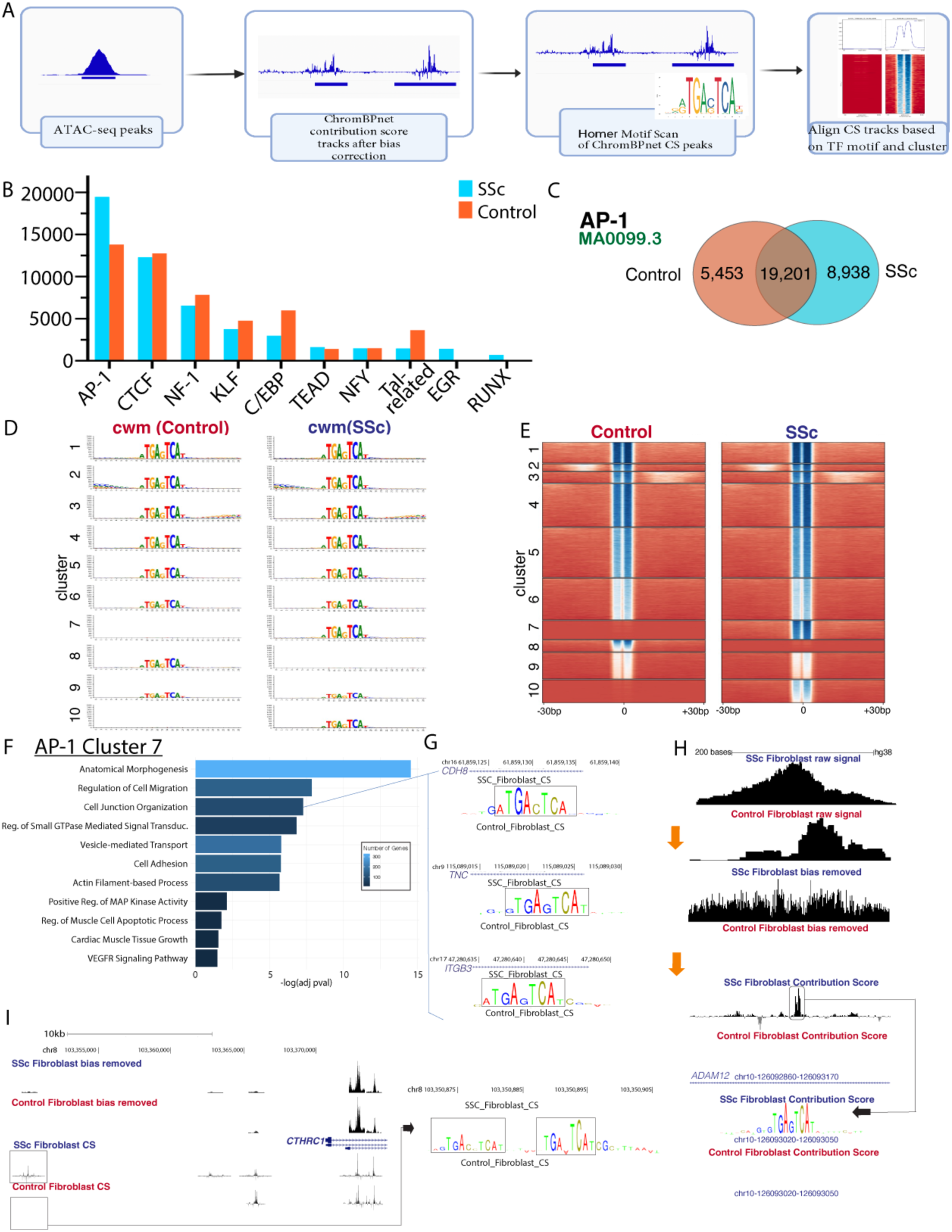
ChromBPNet analysis of 6880 control fibroblast nuclei and 5039 SSc fibroblast nuclei. A- Graphic representation of analysis pipeline for ATAC pseudobulk ChromBPNet analysis B- Number of seqlets detected by transcription factor family (largest number for each family depicted) in SSc and control fibroblasts by TF-MoDISco genome sampling. C- Number of AP-1 seqlets (as detected for the MA0099.3 FOS::JUN matrix profile) unique to and shared by SSc and control fibroblasts. D- Contribution weighted matrices for SSc and control fibroblasts by cluster E- Heatmap of AP-1 contribution score dot product for each seqlet/chromatin region divided by SSc and control fibroblasts into 10 clusters. Darker blue indicates higher contribution score. F- Pathways enriched by gene ontology for the AP-1 cluster 7 chromatin regions. G- Contribution score track depictions for SSc and control fibroblasts of 3 chromatin regions (annotated to *CDH8, TNC,* and *ITGB3*) enriched in the cell junction organization pathway, from AP-1 cluster 7. H- ChromBPNet analysis of *ADAM12* intronic region with AP-1 footprinting for SSc and control fibroblast pseudobulk ATAC data. Multiple levels from pseudobulk raw signal, bias removed signal, and contribution score tracks depicted. I- Pseudobulk bias removed signal and contribution score tracks for *CTHRC1* region with duplicate AP-1 footprinting in SSc fibroblasts.

The most frequently identified de novo seqlet in SSc and control fibroblast peaks matched the consensus AP-1 binding site. Unique and overlapping AP-1 binding sites and their computed contribution score products were identified in both control and SSc-ILD fibroblasts (Figure 3C). We then clustered seqlets and visualized the average contribution weight matrix for each cluster (Figure 3D, 3E, Supp Fig 9A,B). Although scores were higher in nearly every cluster for SSc fibroblasts, cluster 7 had distinctly higher scores in SSc, while cluster 9 had higher scores in control fibroblasts. The AP-1 cluster 7 seqlets were annotated to nearest genes by ChIPseeker, followed by pathway analysis by Gene Ontology (Figure 3F). AP-1 binding was notably stronger in genes like *CDH8, TNC,* and *ITGB3,* involved in Cell junction organization (Figure 3G). The computational process of ChromBPNet in refining the pseudobulk ATAC-seq data is shown in analysis of *ADAM12,* a metalloprotease whose expression is increased in lung (Figure 2D) and skin myofibroblasts^26^, correlates with the severity of skin disease^27^, and predicts response to anti-TGFβ therapy in SSc skin^28^. A strong TF footprint for AP-1 is identified in the *ADAM12* intron 11 in SSc fibroblasts (Figure 3H). Similarly, a duplicate AP-1 site was detected upstream of the transcription start site of the activated myofibroblast marker *CTHRC1,* a significant marker of pathogenic fibroblasts in IPF and murine fibrosis^13,29^ (Figure 3I).

RUNX, EGR, C/EBP, and TEAD motifs were analyzed in an analogous manner. RUNX footprints were present in a small number of seqlets, yet markedly increased in SSc fibroblasts (Supp Fig 10A,11) with enriched pathways linked to genes involved in Multicellular Organism Development and Anatomical Structure Morphogenesis (cluster 9), and Regulation of Signal Transduction and Anatomical Structure Development (cluster 10; Supp Fig 10B-E). Other genes linked to lung fibrosis with upregulated expression and enhanced RUNX binding in SSc fibroblasts included *FHL2, PAPPA* and *ECM2,* as well as *CTTNB2,* a gene previously linked to dermal fibrosis (Supp Fig 11C-F).

EGR clusters 1, 2, 9, and 10 had the greatest contribution score increases in SSc fibroblasts, and were associated with Actin-filament Based Processes and Cell Morphogenesis (Supp Fig 10F-H, Supp Fig 11G,H). EGR footprinting was present in intron 2 of *ADAM12* (Supp Fig 10I), which as mentioned above is also regulated by AP-1 (Figure 3H). TEAD footprints were more evenly distributed to CREs in SSc and control fibroblasts (Supp Fig 12A,B). TEAD cluster 8 contained higher contribution scores in SSc fibroblasts, and was associated with Regulation of Actin-Filament Based Processes and Cardiac Muscle Tissue Growth, consistent with the TEAD TFs known role in muscle differentiation (Supp Fig 12C,D)^30^. C/EBP-binding seqlets were decreased in SSc fibroblasts, most strikingly in cluster 8 (Supp Fig 9C,D).

### Altered Chromatin accessibility of macrophage populations in SSc-ILD

As in prior scRNA-seq studies, we identified macrophage clusters implicated in driving fibrosis in SSc-ILD and IPF by increased expression of *SPP1* and other marker genes (Figures 1A, 4A)^14,31^. We combined the SPP1^hi^ macrophages to examine DARs between SSc-ILD and control lungs, finding 5,194 DARS more accessible in SSc SPP1^hi^ macrophages, and 4,133 in control (Figure 4B, Supp Table 3). Upregulated pathways included Cell-matrix Adhesion, PDGFR Signaling, and ERK1/2 Cascade (Figure 4C). Top DARs in SSc SPP1^hi^ macrophages were annotated to *CHIT3, PSD3,* and PRC (Figure 4E). ChromVAR identified bHLH-ZIP (USF1, USF2, MXI1, MITF, TFEB), EGR and NFAT TFs as highly enriched SSc, while CTCF motifs were enriched in control SPP1^hi^ macrophages (Figure 4D).

**Figure 4.**
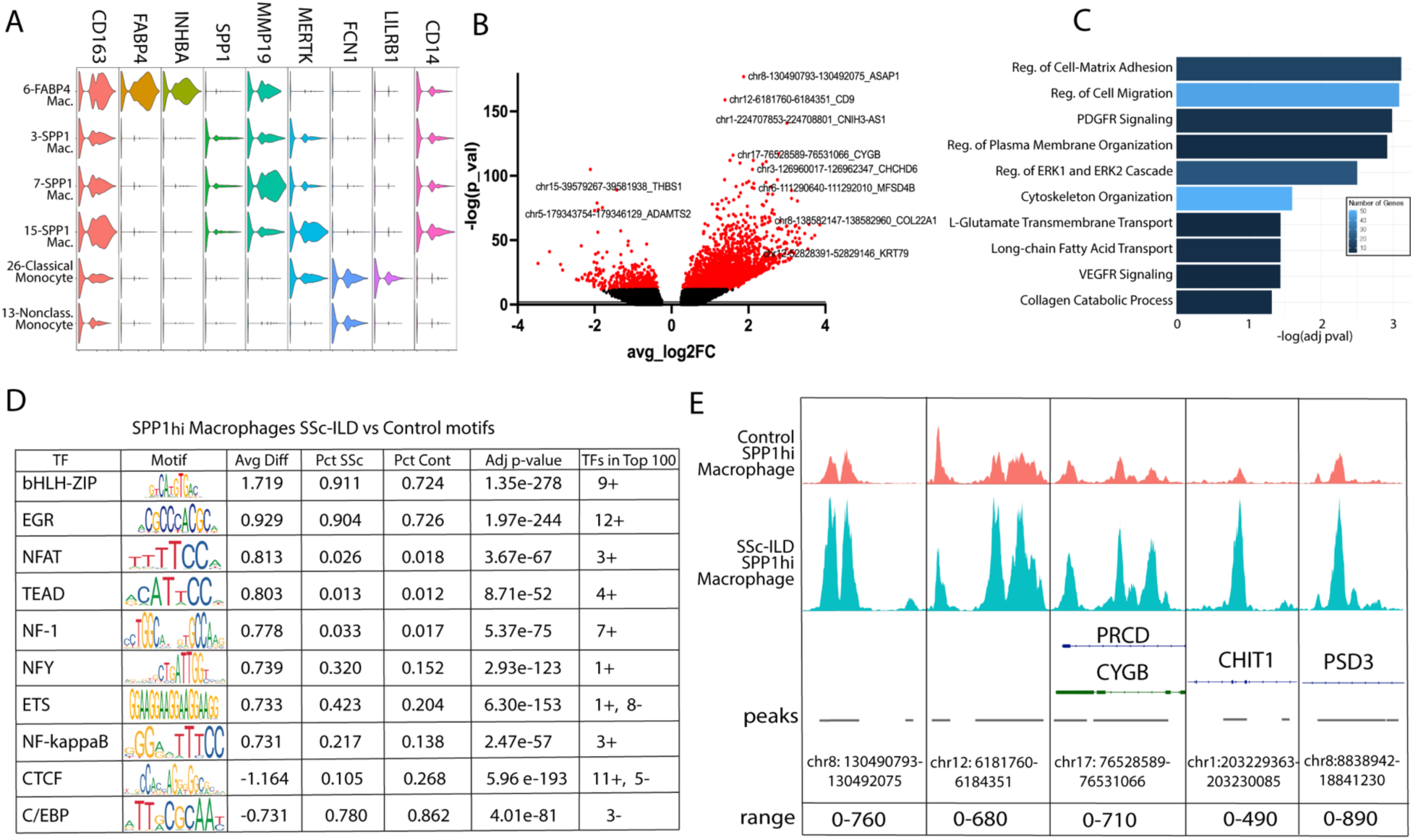
A- Gene expression of myeloid cell type markers B- SSc SPP1^hi^ Macrophages vs Control SPP1^hi^ Macrophages differentially accessible regions with labeling of select DARs and the nearest annotated gene. C- Pathway enrichment analysis of chromatin regions more accessible in SSc vs Control SPP1^hi^ macrophages. D-TF Motifs enriched in SSc SPP1^hi^ Macrophages vs Control SPP1^hi^ Macrophages, ranked by absolute value of average difference and separated into positively and negatively enriched. Final column notes number of TFs in the same class/family present in the top 100. E- Plot of pseudobulk enriched peaks differentially accessible in SSc SPP1^hi^ Macrophages

ChromBPNet analysis predicted bias-corrected TF binding at single base pair resolution from the pseudobulk SPP1^hi^ macrophages. TF-MoDISco indicated that Ets-related binding sites were the most common, followed by C/EBP, AP-1, and CTCF (Figure 5A). AP-1, bHLH-ZIP, and EGR motifs were increased in SSc SPP1^hi^ macrophage seqlets compared to control. In examining AP-1 contribution scores genome wide, cluster 7 had distinctly high scores in SSc (Figure 5B, Supp Fig 16 A,B) and was linked to pathways including Cell-matrix Adhesion and Cell Migration (Figure 5C,D). Cluster 8 had high scores in controls and was enriched for ERBB and JAK-STAT signaling (Figure 5E,F). bHLH-ZIP footprinting for the motif examined (MA0093.3) was mildly increased for unique SSc seqlets (Figure 5G, Supp Fig 16 C,D), with cluster 9 containing higher scores in SSc and linked to Metalloendopeptidase Activity and Autophagy (Figure 5I,J). In SSc, higher bHLH-ZIP contribution scores were present in the *CCL18* promoter, a chemokine ligand influencing collagen production by fibroblasts and highly upregulated by SPP1^hi^ macrophages^32^.

**Figure 5.**
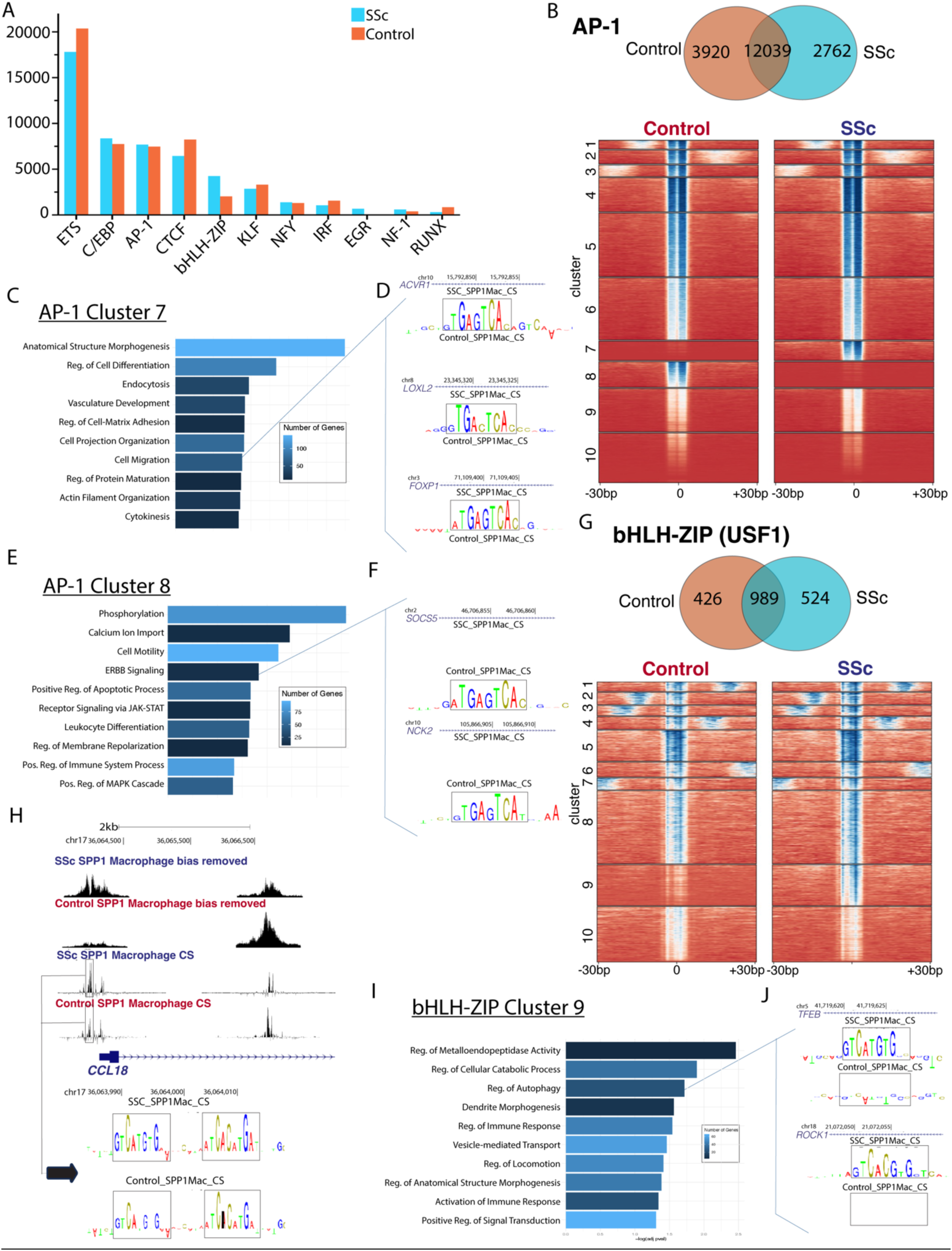
SPP1^hi^ Macrophage ChromBPNet Analysis of 3968 control nuclei and 3902 SSc nuclei. A- Number of seqlets detected by transcription factor family (largest number for each family depicted) in SSc and control SPP1^hi^ Macrophages by TF-MoDISco genome sampling. B- Number of AP-1 seqlets unique to and shared by SSc and control SPP1^hi^ Macrophages. Heatmap of AP-1 contribution score dot product for each seqlet/chromatin region divided by SSc and control SPP1^hi^ Macrophages into 10 clusters. Darker blue indicates higher contribution score. C- Pathways enriched by gene ontology for the AP-1 cluster 7 chromatin regions. D- Contribution score track depictions for SSc and control SPP1^hi^ Macrophages of 3 chromatin regions (annotated to *ACVR1, LOXL2* and *FOXP1*) enriched in the cell migration pathway, from AP-1 cluster 7. E- Pathways enriched by gene ontology for the AP-1 cluster 8 chromatin regions. F- Contribution score track depictions for SSc and control SPP1^hi^ Macrophages of 2 chromatin regions (annotated to *SOCS5* and *NCK2*) enriched in the ERBB signaling pathway, from AP-1 cluster 8. G- Number of bHLH-ZIP seqlets unique to and shared by SSc and control SPP1^hi^ Macrophages. Heatmap of bHLH-ZIP contribution score dot product for each seqlet/chromatin region divided by SSc and control SPP1^hi^ Macrophages into 10 clusters. Darker blue indicates higher contribution score. H- Pseudobulk bias removed signal and contribution score tracks for *CCL18* region with duplicate bHLH-ZIP footprinting in SSc and control SPP1^hi^ Macrophages. I- Pathways enriched by gene ontology for the bHLH-ZIP cluster 9 chromatin regions. J- Contribution score track depictions for SSc and control SPP1^hi^ Macrophages of 2 chromatin regions (annotated to *TFEB* and *ROCK1*) enriched in the regulation of autophagy pathway, from bHLH-ZIP cluster 9.

### Gene regulatory network construction

To further examine the dysregulated gene regulatory networks of fibroblasts and SPP1^hi^ macrophages, we constructed TF-CRE-gene linkages from our multiome RNA/ATAC-seq data (Figure 6A). To better account for the sparsity of single-nucleus genomic data and to utilize multi-omics representations delineating cellular states, we first inferred latent representations using hierarchical causal modeling for single cell multi-omics data (HALO) and constructed metacells of paired RNA-seq and ATAC-seq from our multiome chemistry dataset (n=6 SSc, 7 Control samples)^33^. Metacells are small groups of cells that are more granular than clusters, with each metacell representing a distinct cellular state^34,35^. We then utilized our larger independent scRNA-seq dataset (n=17 SSc, 13 control lungs)^36^ to identify DEGs and CREs (both promoters and more distal elements) occurring within 250 kilobases of the DEGs narrowed to DARs for the comparison of interest. Enriched TF motifs within these DARs were identified through motif matching, followed by TF-CRE-gene linkages.

**Figure 6.**
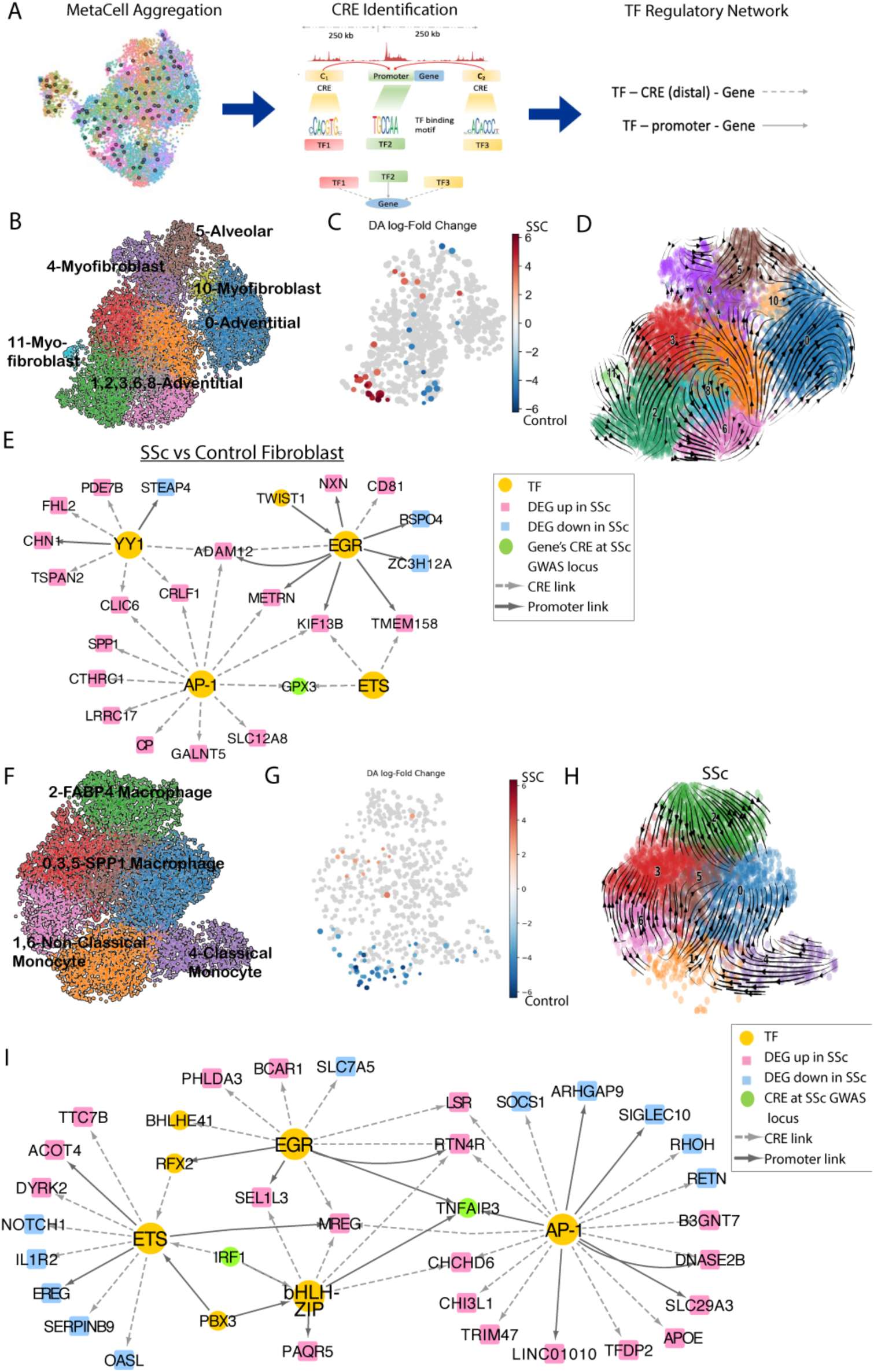
A- Graphic representation of analysis pipeline for multiome mesenchymal and myeloid subsets: 1) Metacell creation using paired ATAC-seq and RNA-seq data to decrease sparsity, 2) Cis-regulatory elements identified by regressing the expression level of a gene of interest (DEGs for the same population comparisons) using accessibility values of peaks located within 250kbp of the transcription start site. Enriched TFs then identified in the differentially accessible CREs. 3)TF regulatory networks constructed by integrating both CRE-gene and CRE-TF relationships. B- Multiome samples fibroblast clustering (smooth muscle, pericytes, and doublet nuclei excluded from this umap) C- Visualization of Milo differential abundance testing results: each point represents a neighborhood, and points are colored according to the log fold change (logFC) in cell abundance between disease and control cells. Red indicates neighborhood enriched in SSc condition and blue indicates neighborhood enriched control condition D- RNA velocity of multiome fibroblasts demonstrating suggested differentiation of myofibroblasts from both alveolar and adventitial subpopulations E- Gene regulatory networks for top enriched TFs (ranked by adj p-value) in SSc fibroblasts vs Control fibroblasts. DEGs for this comparison were generated from a larger scRNA-seq dataset of 17 SSc and 13 control lungs. Only DEGs with abslog2FC >1 are depicted. F- Multiome samples myeloid clustering with cell type identifications G-Visualization of Milo differential abundance testing results. Red indicates neighborhood enriched in SSc condition and blue indicates neighborhood enriched control condition H-RNA velocity of multiome SSc myeloid cells I- Gene regulatory networks for top enriched TFs (ranked by adj p-value) in SSc SPP1^hi^ macrophages vs Control SPP1^hi^ macrophages. DEGs for this comparison were generated from a larger scRNA-seq dataset of 17 SSc and 13 control lungs. Only DEGs with abslog2FC >1 are depicted.

For the fibroblast nuclei we first re-clustered the multiome samples based on HALO representations, followed by cell annotation, construction of metacells, and identification of cell states enriched in SSc fibroblasts using neighborhood-level differential abundance testing (Figure 6B,C, Supp Fig 18A)^37^. RNA velocity was applied to assess cellular dynamics of the myofibroblast population^38^. Velocity projections indicated that myofibroblasts arise from both alveolar and adventitial fibroblast populations (Figure 6D). In comparing SSc vs control fibroblasts, AP-1, EGR, YY1, and ETS-related family TFs had the largest positive average difference in motif activity (Figure 6E). The TF ZNF148 had the greatest number of combined proximal and distal CRE links (551) to DEGs, with IRF1 and KLF4 also having greater than 500 links. For this comparison, 65 TFs showed statistically significant differences in both gene expression and motif activity, 28 had discordant directionality for these differential values while 4 TFs were downregulated with negative motif activity differences, and 33 TFs were upregulated with positive motif activity differences (Supp Table 4). The AP-1 TFs JUND and JDP2 had a top 5 positive change for both measures. Members of the AP-1 family with significantly enriched motif activity (JUND, JUNB, JDP2, FOSL1) were linked to 280 unique DEGs (243 upregulated, 37 downregulated) for the SSc vs control fibroblasts, including the upregulated genes *CTHRC1, ADAM12, IGFBP2, SPP1* and *TGFBI.* Enriched EGR TFs EGR1, EGR2, and EGR3 were linked to 320 unique DEGs (273 upregulated, 46 downregulated) including the upregulated genes *ADAM12, TMEM158, IGFBP2,* and *SPP1*.

Myeloid cells were examined in a similar manner. SPP1^hi^ macrophages were increased in SSc, while non-classical monocytes were increased in control samples (Figure 6F,G, Supp Fig 18D). Velocity projections predicted that SPP1^hi^ macrophages arise from both the FABP4^hi^ macrophage and monocyte populations, although a larger transition from monocytes to SPP1^hi^ macrophages was indicated in fibrotic lungs. (Figure 6H, Supp Fig 18F). In examining motif activity for this model, EGR, AP-1, bHLH-ZIP, and Ets-related TFs had the largest positive average difference in motif activity (Figure 6I). The TF ZNF148 had the greatest number of combined proximal and distal CRE links (146) to DEGs. For this comparison, 29 TFs showed statistically significant differences in both gene expression and motif activity, 12 had discordant directionality for these differential values while 7 TFs were downregulated with negative motif activity difference, and 10 TFs were upregulated with positive motif activity difference (Supp Table 5). TFEB and MXI1 had a top 5 positive change for both measures. Members of the bHLH-ZIP family with significantly enriched motif activity (MITF, MXI1, and TFEB) were linked to 64 unique DEGs (53 upregulated, 11 downregulated) for the SSc vs control SPP1^hi^ macrophage population, including the upregulated genes *PAQR5, MREG, SEL1L3, RTN4R,* and *APOE.* Enriched AP-1 TFs FOSL1 and JDP2 were similarly linked to 69 unique DEGs (54 upregulated, 15 downregulated) including the upregulated genes *CHIT1, CHI3L1, MMP7, DNASE2B, RTN4R,* and *CDH23*.

## Discussion

Here, we delineate dynamic chromatin changes and altered TF binding in fibrotic lung disease. Our study provides a comprehensive map of TF regulation in the human lung and focuses on primary TFs regulating the transition of fibroblasts and macrophages to profibrotic states^36,39^. Our study generated the largest multiomic lung cell atlas of both control and fibrotic ILD. Applying ChromVAR we inferred the TFs regulating cell states in healthy and fibrotic lungs. Applying the deep learning model ChromBPNet, we detect TF binding at single nucleotide resolution, confirming known TF motifs and inferring the altered binding in profibrotic fibroblasts and macrophages in an unbiased manner. HALO metacells enabled robust trajectory analysis and linked TF and CREs to regulated genes, supporting the TF binding indicated by ChromBPNet.

In SSc fibroblasts, we observed enriched binding to AP-1 TF motifs and high contribution score footprints of AP-1 binding in multiple fibrosis-related genes, such as collagen. AP-1 TFs, comprised of homo- and hetero-dimers of Jun, Fos, Atf, and Maf, are implicated in fibrosis^40,41^. Fra-2 transgenic mice develop skin fibrosis, ILD, and vascular disease^42^. AP-1 is upregulated in a TGF-β-dependent manner in SSc dermal fibrosis, with reduction in collagen synthesis *in vitro* and prevention of dermal fibrosis *in vivo* following treatment with a selective AP-1 inhibitor^43^. Our data suggest AP-1 regulates key myofibroblast marker genes, including *CTHRC1* and *ADAM12. CTHRC1* has been identified as a myofibroblast marker in fibrotic lung diseases ranging from IPF to COVID-19^13,44^. Its deletion either aggregates or ameliorate fibrosis in lungs and liver respectively. Deletion of *ADAM12* blocks the development of profibrotic progenitors^45^.

The significant redistribution of accessible AP-1 sites across the genome, and the identification of clusters of seqlets with AP-1 motifs showing no change or increased binding in control fibroblasts highlights the complex action of this TF family. Several mechanisms are likely in play. Specific AP-1 TFs may increase accessibility, while others decrease accessibility in SSc-ILD fibroblasts. For example, increased JunB replacing Jun regulates an enhancer upstream of COL1A2^40^. Although binding of other TFs may influence binding to AP-1 sites locally or distally, our data remarkably did not indicate binding of TFs immediately adjacent to the AP-1 sites, which appear rather as islands of enhancer binding. Distal TF interactions, DNA loop formation, and/or epigenetic changes at the level of histone or DNA modifications appear more likely explanations. Determining the molecular mechanisms driving AP-1 binding differences between healthy and SSc-ILD fibroblasts will clarify determinants of AP-1 action, the resulting gene dysregulation, and strategies for intervention.

Although RUNX binding was inferred in fewer seqlets than many TFs, its footprinting was more consistently increased in SSc and appeared in few control fibroblast OCRs. RUNX proteins bind DNA after complexing with their coactivator core-binding factor β-subunit(CBFB)^46^. Inhibition of RUNX1 *in vitro* blocks myofibroblast differentiation^47^, RUNX footprinting is enriched in SSc dermal fibroblasts^48^ and RUNX1 has also been implicated in myofibroblast differentiation in scRNA-seq studies of SSc skin^26^. Only one of 16 genes highlighted in cultured SSc dermal fibroblast ATAC-seq analyses was identified in our analysis: CTTNBP2^48^. This may reflect differences between lung and skin myofibroblasts or the *in vitro* versus *in vivo* analysis enabled by snATAC-seq. Although pathway analysis of genes proximal to RUNX seqlets showing increased binding in SSc-ILD did not suggest traditional fibrosis pathways, seqlets associated with genes showing increased expression in SSc-ILD myofibroblasts included genes implicated in lung fibrosis. Notably, FHL2 is one of 12 genes overexpressed in both rat and human lung fibrosis^49^. It binds to focal adhesions and transmits mechanical signals to regulate gene expression^50^ and its pharmacologic inhibition by Arbutin ameliorates rat lung fibrosis^51^. ECM2 and PAPPA are markers of IPF and IPF severity^52^. Senescent cells secrete increased PAPP-A, suggesting a role in IPF^52,53^. ECM2 deletion inhibits skeletal muscle-derived satellite cell differentiation^54^, suggesting a possible role in myofibroblast differentiation. Further studies will be needed to clarify how RUNX proteins control myofibroblast differentiation. However, the availability of small molecule inhibitors for both RUNX-DNA binding and CBFB-RUNX binding^55–57^, as well as its significant post-transcriptional regulation by ubiquitin-proteasome degradation^58^, offers a variety of approaches to inhibiting RUNX activity.

EGR1 is implicated in fibrotic signals from IL-13 and TGF-β ^59,60^. EGR1 deletion inhibits wound healing and myofibroblast differentiation, while overexpression enhances wound repair^61^ and upregulates genes associated with wound healing and fibrosis^62^. EGR1 deletion also protects against renal fibrosis^63^, but aggravates experimental liver fibrosis^64^. EGR2 and EGR3 both mediate TGF-β dependent signals including collagen expression and myofibroblast differentiation but regulate distinct genes^65,66^. Previous *in vitro* studies indicate SSc dermal fibroblasts have enhanced EGR1 accessibility^66^, suggesting EGR1 may play parallel roles in myofibroblast differentiation in the skin and lungs.

In SPP1^hi^ macrophages, we identified enrichment of bHLH-ZIP, AP-1, and EGR motifs in SSc compared to controls, as well as high contribution scores of these TFs. The bHLH-ZIP TFs demonstrated the most significant motif enrichment, and the largest increase in SSc seqlets when identified *de novo* by ChromBPNet and TF-MoDISco, suggesting a distinct regulatory program in SPP1^hi^ macrophages compared to fibroblasts. The bHLH-ZIP TFs include USF1 and USF2, as well as MiT/TFE members MITF, TFEB, MXI1, and TFE3^67^. Because these TFs share the same consensus motif, we cannot clearly distinguish which is most important. TFEB is a master regulator of lysosomal biogenesis and autophagy^68^, while members also act in mitophagy, lipid catabolism, and tumor-associated macrophage polarization^69–71^. They link mitochondrial and lysosomal functions, both implicated in SSc^72,73^. We previously identified TFEB as a putative regulator of pro-fibrotic macrophages in SSc-ILD^36^. USF1 is linked to aging-related macrophage function decline^74^ and liver fibrosis^75^. The enriched duplicate bHLH-ZIP footprint in proximal CCL18 is notable, as elevated CCL18 is associated with SSc-ILD severity and performs well as a diagnostic biomarker of SSc-ILD^76,77^. CCL18 is decreased in the serum and skin biopsies of patients with SSc following IL-6 inhibition^78^, and is downregulated by JAK inhibitors^79^. CCL18 increases collagen production by lung fibroblasts^80^, and genetic variability in *CCL18* affects gene expression and mortality in IPF^81^. bHLH-ZIP TFs may represent a key influence on the feedback loop between fibrotic macrophages and fibroblasts, warranting further investigation.

To our knowledge, this is the first large snATAC-sequencing study of adult healthy and fibrotic lungs, providing a rich resource on chromatin accessibility in the human lung. We utilized deep learning models to uncover intricate sequence rules governing TF binding, providing insights into transcriptional programs regulated by CREs in fibroblasts and macrophages. As our study used explanted tissue from patients with severe SSc-ILD undergoing lung transplants, it may not reflect changes in mild disease or the broader SSc-ILD population. As OCRs were annotated by least base-pairs distance, not all annotations may reflect the regulated gene, as CREs can regulate distant genes via long-range interactions. While our study focused on OCRs and TF binding sites, additional non-coding regulatory mechanisms including long non-coding RNAs^82^, microRNAs^83^, and transposable elements^84^ may also influence gene expression.

In summary, utilizing multiomic single-cell approaches in advanced SSc-ILD and control lungs, we identified cell populations with significant changes in chromatin accessibility, altered TF activity in SSc-ILD fibroblasts and SPP1^hi^ macrophages, and the resulting TF-CRE-gene regulatory networks. The snATAC-seq data maps TFs determining cell populations in healthy and fibrotic lungs, highlighting cis-regulatory changes responsible for the diseased phenotypes of profibrotic macrophages and myofibroblasts. Our study lays the groundwork for functional validation of these differential CREs, their target genes, and the TFs driving them. This knowledge could guide the development of targeted therapeutics aimed at modulating TF or their coregulators to mitigate progressive fibrotic lung disease.

## Methods

### snATAC-seq/multiome Analysis

The University of Pittsburgh Institutional Review Board approved procedures involving human samples. Explanted subpleural peripheral lung tissue was digested to single-cell suspensions using Liberase DL (Roche) and bovine pancreas DNAse I (Sigma-Aldrich) as previously described^15^. Samples were processed immediately upon receiving from the operating room. Some samples were partially enriched for fibroblasts using magnetic-activated depletion of CD45, CD31, CD163, and/or CD326 positive cells, with 50/50 loading of the enriched and non-enriched cell suspension at the time of running snATAC-seq as detailed in Supplementary Table 1. To increase data quality dead cell removal (STEMCELL, Annexin V kit) and CD66 depletion were added to sample processing for all digestions as well starting with sample SC421. For the snATAC-seq only samples, single-cell suspensions were split, with a portion used for performing scRNA-seq as previously described^14^, and the remaining suspension used for nuclei generation and snATAC-seq as previously described^85^. For the multiome samples, transposition of nuclei and library preparation were carried out per the Chromium Next GEM Single Cell Multiome ATAC + Gene Expression kit protocol (10x Genomics Catalog no: 1000283). Libraries were sequenced using the Illumina NovaSeq 6000 platform.

CellRanger ATAC pipeline (v2.0.0) and CellRanger Arc pipeline (v2.0.2) with h38 human genome reference were used for initial read processing, library quality control, peak calling and cell calling. Samples were filtered for: minimum features>200 and <150,000, peak region fragments>200 and < 150,000, nucleosome signal<10. For the snATAC-seq samples, the R package Signac (v1.9.0)^19^ was used for merging samples, unsupervised clustering, and downstream analyses. A unified peak set was created and quantified in each dataset, followed by new fragment objects, Seurat objects, and merging of the 5 samples. Batch correction at the individual sample level was performed using the R package harmony (v0.1.1)^20^. Multiome chemistry samples were aggregated using CellRanger Arc ‘aggr’ for creation of a unified fragment file, followed by disaggregation, filtering, peak calling, and clustering by weighted nearest neighbor analysis. The multiome object peaks were quantified within the snATAC-seq object, followed by merging of these objects. Term frequency-inverse document frequency (TF-IDF) normalization and singular value decomposition (SVD) were performed on the DNA accessibility data using Signac’s ‘RunTFIDF’ and ‘RunSVD’ functions, followed by removal of low-quality nuclei clusters. Peak calling was performed by cluster using MACS2^86^, with this peak set used for downstream analysis and subclustering of the mesenchymal populations. For manual annotation of principal cell types, the UMAP projection of nuclei was queried for clusters showing accessibility of known cell type-specific genes, as well as by gene expression profiles. DARs were calculated by logistic regression test with number of peak region fragments as the latent variable and requiring peak accessibility in 5 percent or more of nuclei in at least one population. Wilcoxon rank sum test with Bonferroni FDR correction was used for DEG testing. Signac was used for display of pseudo-bulk coverage tracks of Tn5 integration for designated cell populations.

For re-clustering of mesenchymal clusters, we subsetted the following clusters: 0,9,17,22,23,31,33, and 34, and performed normalization, dimensionality reduction, batch effect removal, clustering and cell type annotation as described above. Low-quality clusters and clusters expressing gene markers for 2 or more cell types were excluded from downstream analysis. Gene expression data was log-normalized and scaled using Seurat’s ‘NormalizeData’ and ‘ScaleData’ functions. Principal component analysis (PCA) was performed with Seurat’s ‘RunPCA’ function using variable features determined with Seurat’s ‘FindVariableFeatures’ function. Motif activity scores were calculated using chromVAR^22^. To calculate differential motif accessibility between populations a logistic regression test was used with number of peak region fragments and MACS2 peak fragments as latent variables. Chromatin regions were annotated by ChIPseeker^21^, with pathway enrichment performed by GeneOntology enrichment analysis^87,88^, utilizing genes or annotated genes with adjusted p-value<0.05 and absolute log2 fold-change greater than 0.5.

Differentially expressed genes utilized in the construction of DEG-CRE-TF linkages for SSc versus control fibroblast and SSc versus control SPP1hi macrophages were obtained from our previously published dataset of 17 SSc-ILD and 13 control lung scRNA-seq samples^36^.

### ChromBPNet Analysis

#### Pseudobulk snATAC sequencing file generation

Fragment file containing the position of Tn5 integration site along with the cell barcodes for the relevant cell subpopulations were extracted using Signac. Single-nucleus ATAC sequencing BAM files for fibroblasts and SPP1hi macrophages were subsetted from the entire single-cell Multiome and ATAC-seq datasets using Sinto’s ‘filterbarcodes -b’ function on the fragment file (https://timoast.github.io/sinto). Peak files were generated on the bam file using MACS2.

ChromBPNet takes the binary alignment matrix (BAM) files as input along with the peaks called from the pseudobulk BAM file and generates three important outputs. The first is the learned bias pattern from the inaccessible chromatin regions. Tn5 cleavage patterns in these regions are caused by enzyme cutting DNA sequence preferences. The second is the bias-corrected chromatin accessibility tracks that are generated after removing the learned bias from the accessibility signals. The third is, the contribution score which assigns an importance score for each base pair in the open chromatin regions. The score prioritizes base pairs that are predicted to contribute to the accessibility of the region where they are located and in designated cellular context.

Pseudo-bulk ATAC-seq profiles for SSc and control fibroblasts and corresponding SSP1 hi macrophages were used to train bias models on non-accessible genomic regions which were frozen and utilized for training secondary models on accessible (peak) genomic regions. After removing bias, corrected signal profiles for each of the four cell states were used to generate single-base pair resolution contribution scores. TF-MoDISco^23,24^ enabled de novo motif discovery from a sampling of 50,000 seqlets from each cell state which were annotated with JASPAR TF motifs. To complement TF-MoDISco, analysis peaks (30bp maximum) were called using the ChromBPNet contribution score tracks. Sequence contribution scores were visualized by dynseq tracks in the UCSC genome browser^89^. The contribution score peaks were intersected with JASPAR motif coordinates^25^ to enable comprehensive analyses of known TF motifs with high contribution scores occurring within accessible chromatin regions. Homer and Deeptool were used to scan for specific TFs motif in the genome using its PWM. The aligned genomic coordinates for a given TF motif were used to generate cluster-specific aggregate contribution scores. K-Means sub clustering enabled genomic regions to be identified with dynamic aggregate contribution scores in control vs. SSc fibroblasts and in control vs. SSc SPP1^hi^ macrophages. These regions were associated with nearby genes using ChIPseeker.

### Multi-Omics METAcell Construction

We utilized HALO, a variational autoencoder model^33^, to infer latent representations from scRNA-seq and snATAC-seq data. Using the representations, a neighborhood graph was constructed using Scanpy (v1.9.3) to facilitate graph clustering and UMAP embedding visualization^90^. Given the sparse nature of single-cell data, we aggregated individual cells exhibiting similar states into METAcells using COEM framework^91^. These METAcells were constructed based on cell-cell similarities inferred from the latent representations, which accounts for potential time-lag events between RNA-seq and ATAC-seq data.

### Construction of TF-CRE-Gene Linkages

To identify cis-regulatory elements (CREs) from single-cell multiome data, we adopted the nonlinear regression-based framework, DIRECT-NET^92^. Specifically, we employed the nonlinear predictive model XGBoost to regress the expression level of a gene against the accessibility values of candidate distal CREs (defined as peaks within 250kbp upstream and downstream of the transcription start site). CREs were then selected based on the importance scores derived from the XGBoost model. CREs with the importance scores higher than the maximum of the median of importance scores and 0.001 were included for TF regulatory network construction.

To construct a disease-associated TF-gene regulatory network, we first identified differentially expressed genes (DEGs) (logFC > 0.5 and p.adj < 0.05) across conditions (control vs. disease, or comparisons among cell states). We then identified differentially accessible CREs (logFC > 0.1 and p.adj < 0.05) of these DEGs, ensuring that these differentially accessible CREs also had high importance scores. TFs in the differentially accessible CREs and promoter regions were identified using the JASPAR2020 motif database and the motifmatchr function in the ChromVAR package^22,93^. Finally, we constructed a TF-gene regulatory network by integrating CRE-TF with CRE-gene relationships. The resultant network includes two types of regulatory linkages: TF-CRE-gene and TF-promoter-gene. If a SSc GWAS locus is located at CREs of DEGs, this information was overlaid onto the network [GWAS input: https://www.ebi.ac.uk/gwas/efotraits/EFO_0000717]. The network was visualized using Cytoscape software^94^.

For RNA velocity trajectory analysis, the spliced and un-spliced RNA count matrices were generated using Cell Ranger output BAM files with Velocyto CLI(v0.17.17)^38^. These matrices were then fed into scVelo (v 0.3.1)^95^ to compute RNA velocity with ‘Dynamical’ mode.

We evaluated differences in cell abundances associated with disease using the Milo framework for differential abundance testing^37^, with its Python implementation, Milopy (https://github.com/emdann/milopy). Milo analysis was performed using the standard workflow, the KNN graph was generated from the latent representation of scRNA-seq and snATAC-seq data.

## Acknowledgements

The authors would like to acknowledge the Center for Organ Recovery & Education (CORE) as well as the organ donors and their families for the generous donation of tissues used in this study. This research was supported in part by the University of Pittsburgh Center for Research Computing, RRID:SCR_022735, through the resources provided. Specifically, this work used the HTC cluster, which is supported by NIH award number S10OD028483.

## Financial Support

Support for the studies was provided by NIH NIAMS P50AR080612 to RL and HS, DOD SL210018 to RL and HS, NIH NHLBI K08 HL161258 to EV, and from the Pulmonary Fibrosis Foundation, Francis Family Foundation, and National Scleroderma Foundation to EV.

## Conflict of Interest

RL reports grants from Bristol Meyers Squibb, Formation, Moderna, Regeneron and Pfizer, EV reports grans from Boehringer Ingelheim. RL served or serves as a consultant with Abbvie, Mediar, Bristol Meyers Squibb, Formation, Thirona Bio, Sanofi, Boehringer Ingelheim, Merck, Genentech/Roche, EMD Serono, Morphic, Third Rock Ventures, Bain Capital and Zag Bio. RL sits on independent data safety monitoring committees for Advarra/GSK and Genentech. RL holds stock in Thirona Bio Inc and is president and holds stock in Modumac Therapeutics Inc.

## Notes

### Competing Interest Statement

RL reports grants from Bristol Meyers Squibb, Formation, Moderna, Regeneron and Pfizer. EV reports grants from Boehringer Ingelheim. RL served or serves as a consultant with Abbvie, Mediar, Bristol Meyers Squibb, Formation, Thirona Bio, Sanofi, Boehringer Ingelheim, Merck, Genentech/Roche, EMD Serono, Morphic, Third Rock Ventures, Bain Capital and Zag Bio. RL sits on independent data safety monitoring committees for Advarra/GSK and Genentech. RL holds stock in Thirona Bio Inc and is president and holds stock in Modumac Therapeutics Inc.

