## Supplemental Figures for "Altered AP-1, RUNX and EGR chromatin dynamics drive fibrotic lung disease"

### Extended Data File

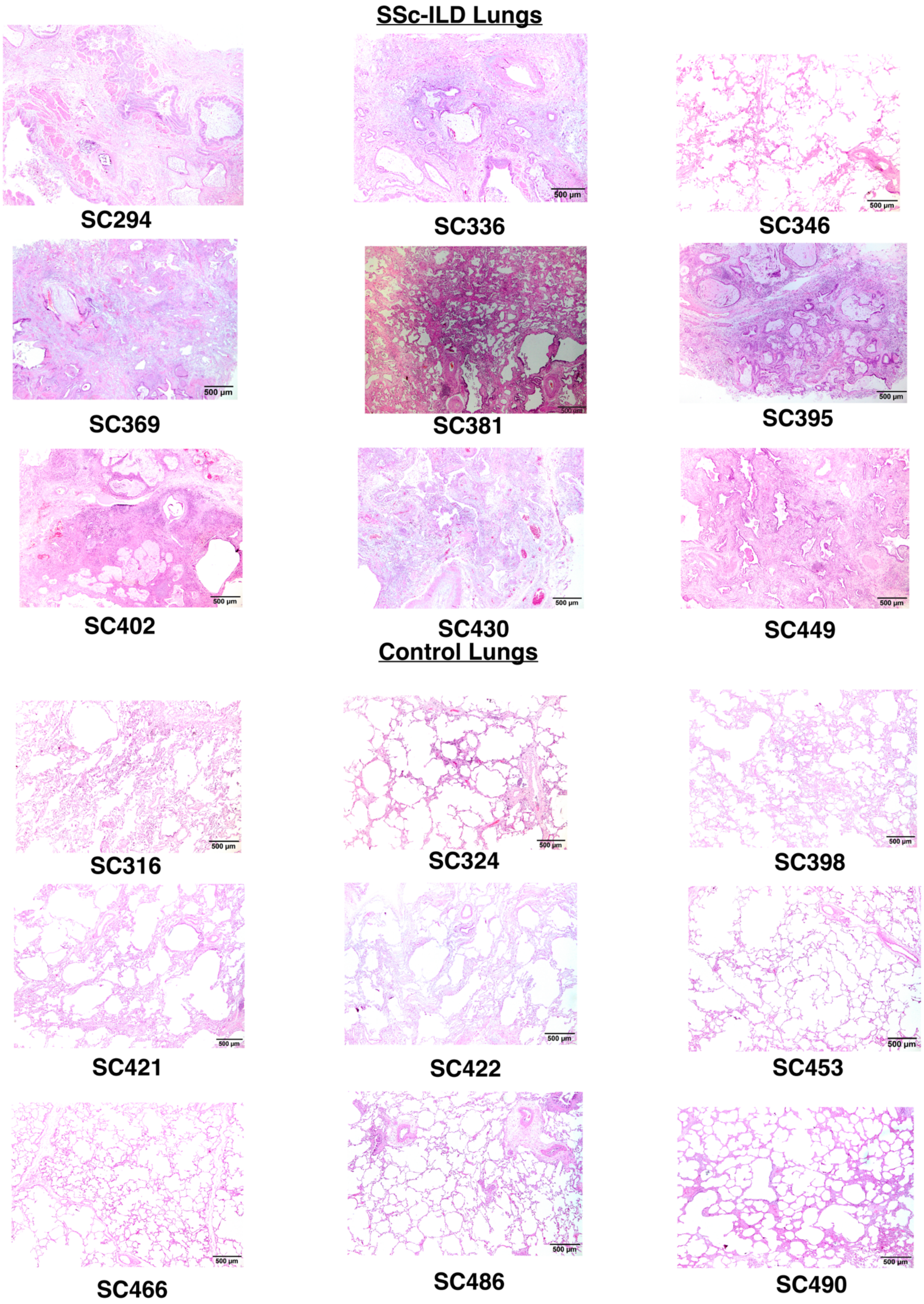


**Supplemental Figure 1**

Histology of adjacent lung tissue for all samples utilized.

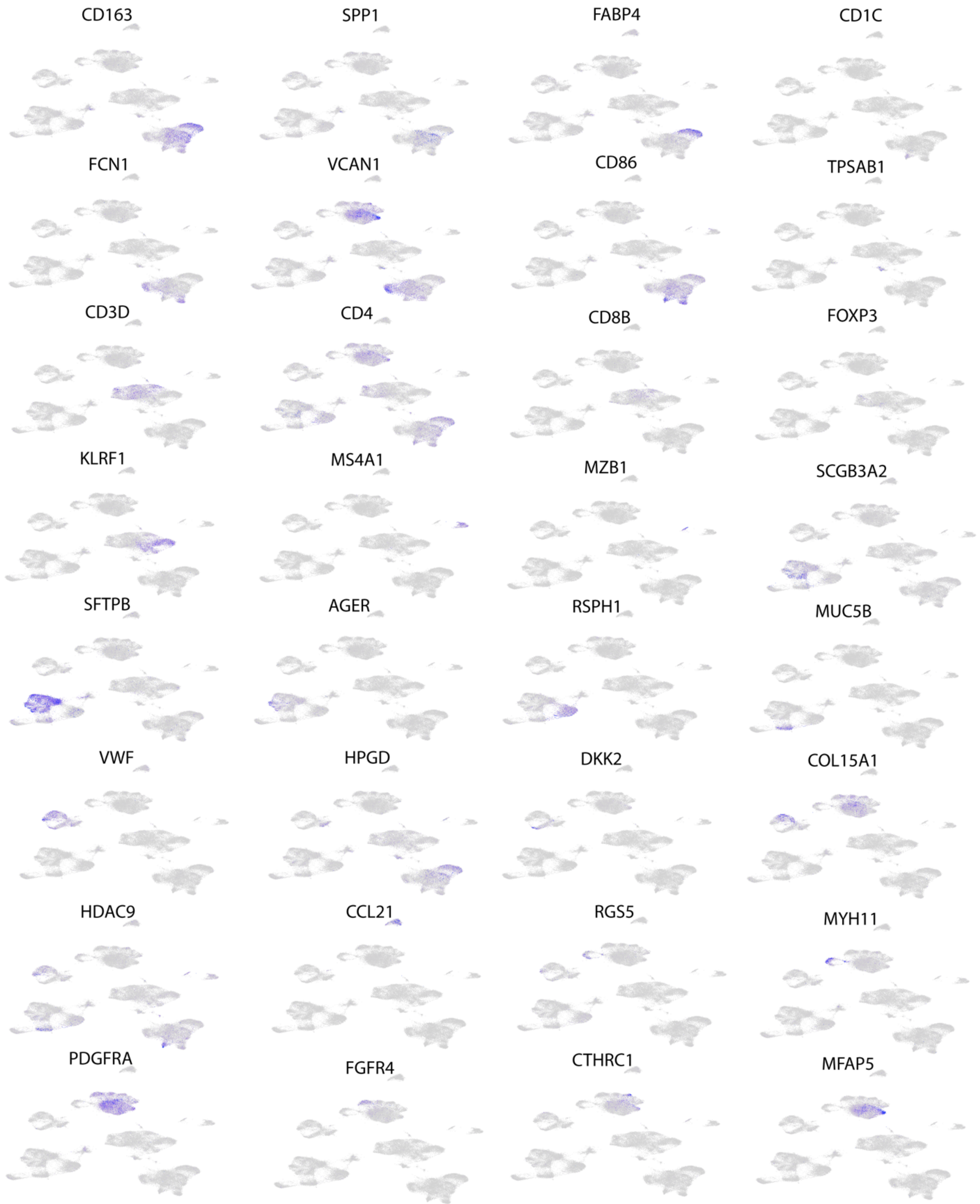


**Supplemental Figure 2**

Gene expression of select cell type markers in clustering of the 9 SSc and 9 Control samples.

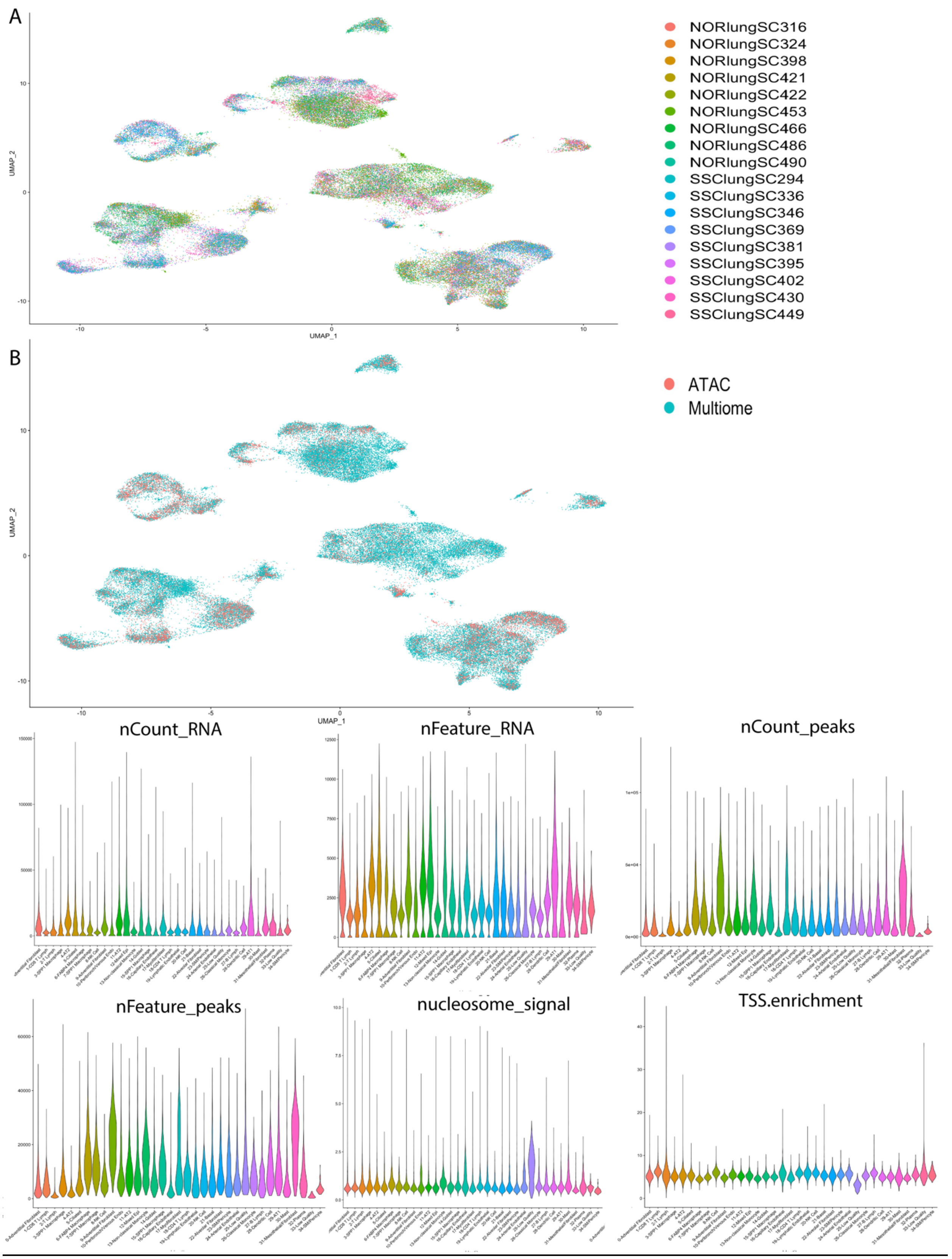


**Supplemental Figure 3**

A-Clustering depicted by individual sample

B-Clustering depicted by chemistry utilized

C-Quality metrics including nCount_RNA, nFeature_RNA, nCount_peaks, nFeature_peaks, nucleosome signal, and transcription start site enrichment per cluster

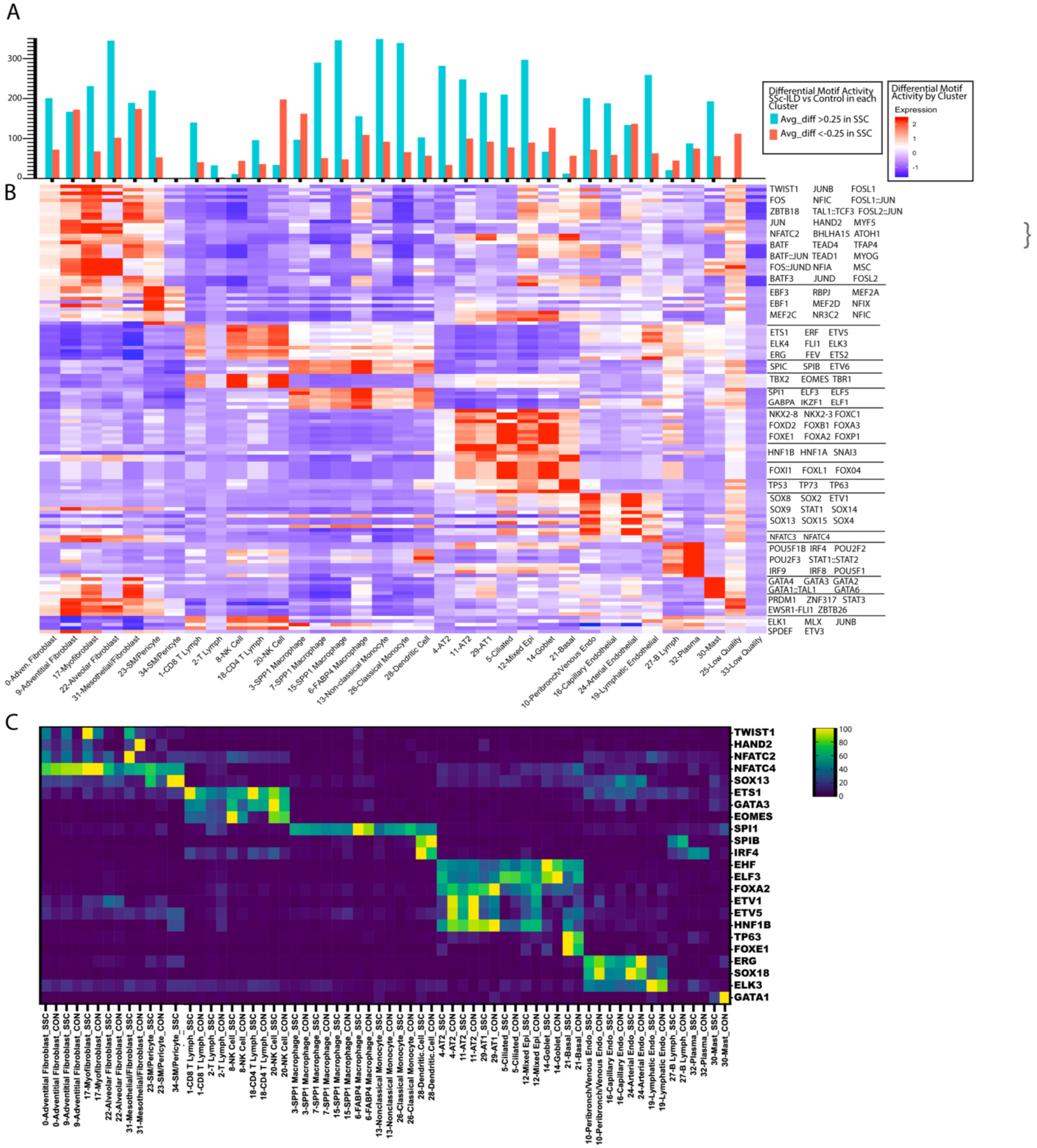


#### Supplemental Figure 4

A- Number of statistically significant (adj p-val<0.05) enriched motifs in SSc versus Control nuclei within each cluster

B- Heatmap of top enriched motifs for each cluster in the clustering of SSc and Control lungs

C- Normalized Gene Expression of transcription factors with cell-type specific expression

A

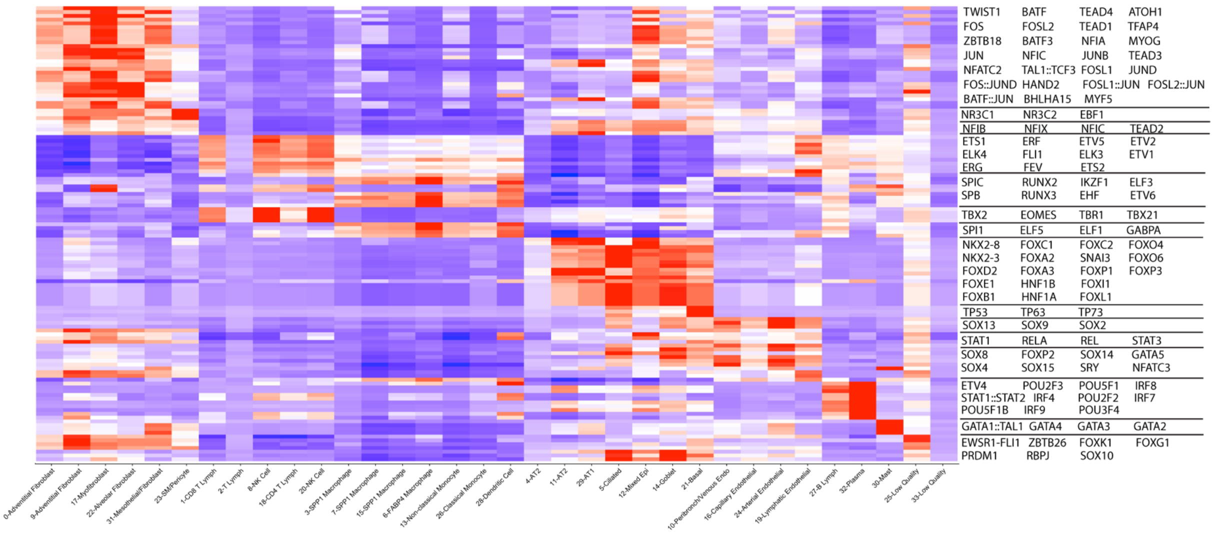


**Supplemental Figure 5**

Heatmap of top enriched motifs by cluster in control lungs only

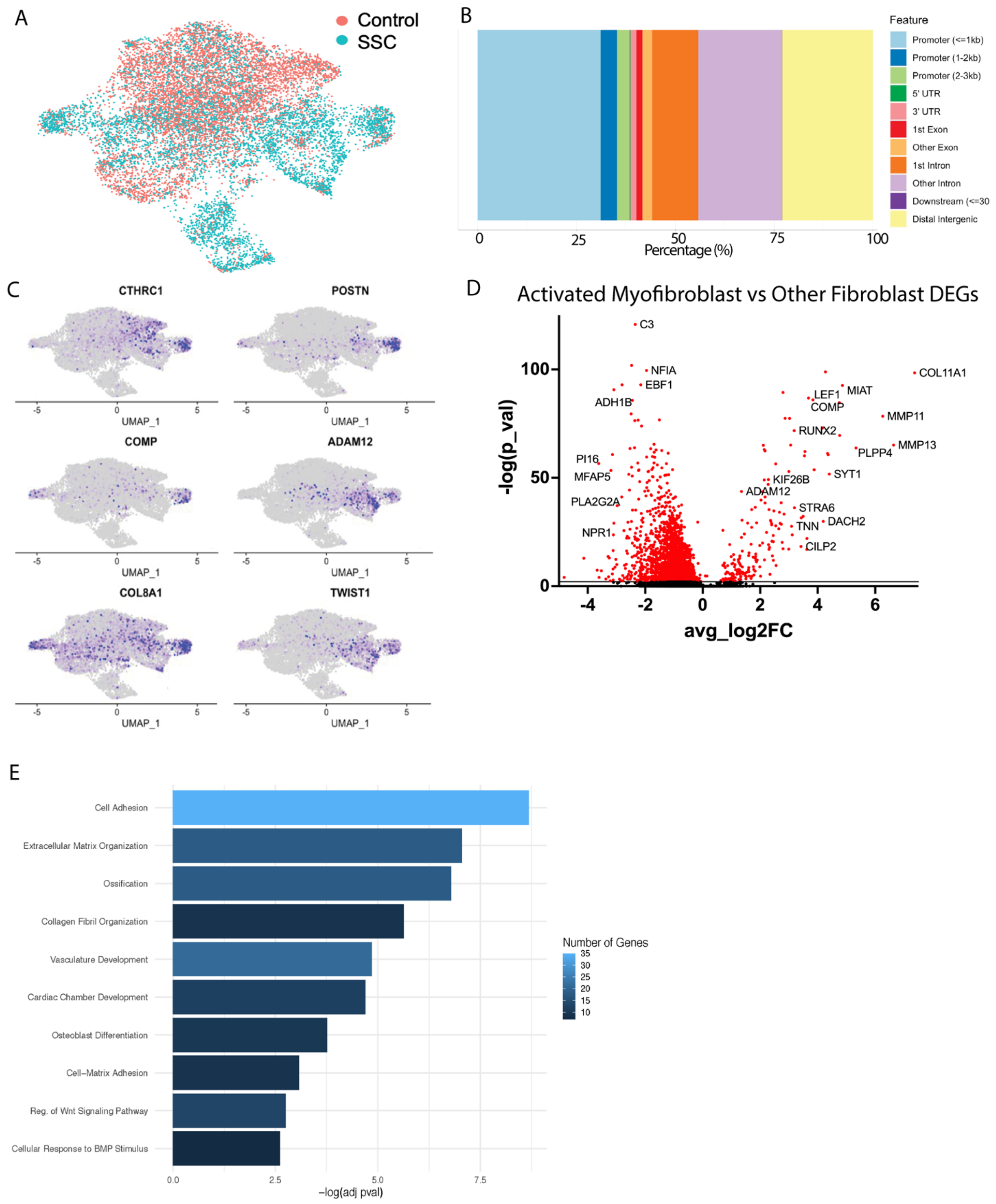


**Supplemental Figure 6**

A- Mesenchymal clustering divided by SSc versus Control origin

B- Annotation of peak regions for mesenchymal clustering. Peaks called per cluster by macs2 with annotation by ChIPseeker

C-Gene expression in mesenchymal clustering of genes enhanced in activated myofibroblasts

D-Differentially expressed genes in activated myofibroblasts vs other fibroblast clusters from multiome samples. Statistically significant genes depicted in red.

E-Pathway enrichment analysis of differentially expressed genes in activated myofibroblasts vs other fibroblast clusters.

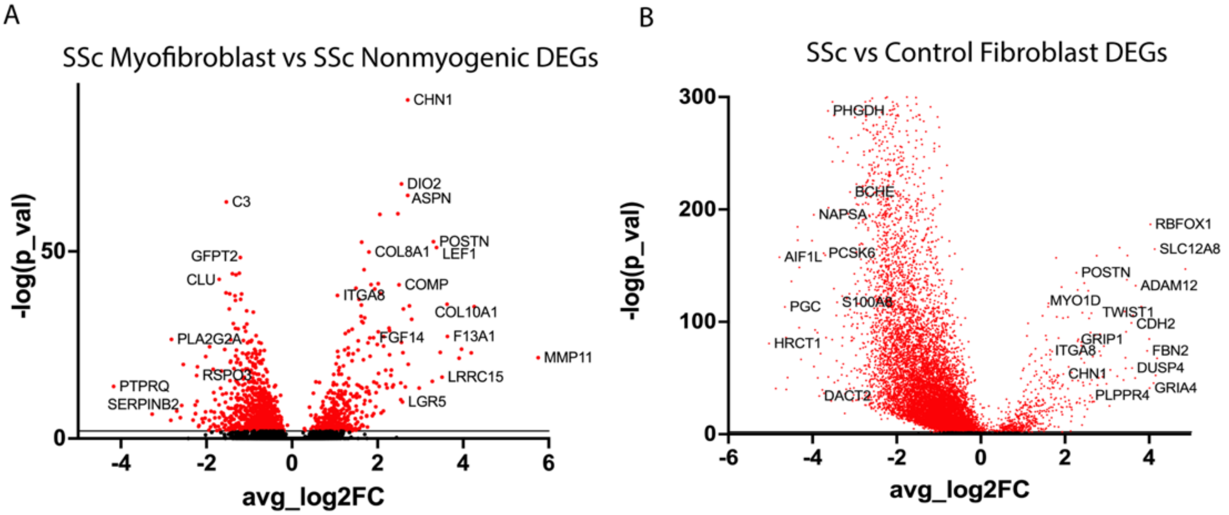


C

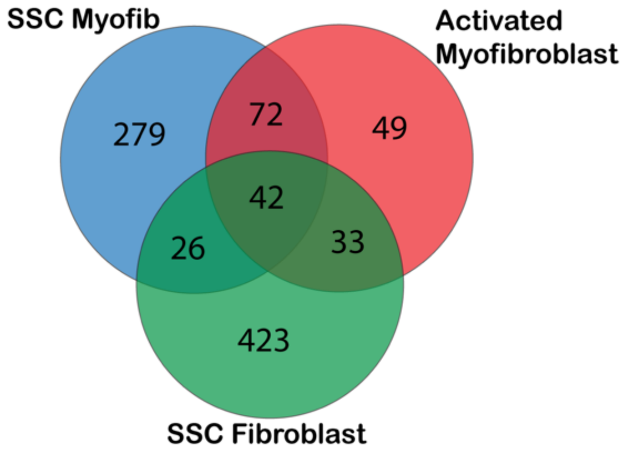


D

Shared DEGs of SSc Fibroblasts, SSc Myofibroblasts, and Activated Myofibroblasts

| | | |
|---|---|---|
| ADAMTS14 | GLIS1 | PDGFC |
| BMP5 | HDAC9 | POSTN |
| CDC42EP3 | HECW1 | PPFIA2 |
| CERS6 | INHBA | PRAG1 |
| CHN1 | ITGA8 | PRR16 |
| COL13A1 | KIAA1211 | ST6GALNAC5 |
| COL24A1 | KIF26B | SUGCT |
| COL8A1 | LEF1 | TGFBI |
| COMP | LINC00511 | TNFRSF19 |
| CSRNP3 | LINC00578 | TSPAN11 |
| DOCK10 | MIR181A1HG | TUSC3 |
| EDIL3 | MRVI1 | TWIST1 |
| EDNRA | NALCN | |
| EEPD1 | NXN | |
| ENTPD1 | PCDH7 | |

**Supplemental Figure 7**

A- Differentially expressed genes in SSc myofibroblasts vs SSc non-myogenic fibroblasts. Statistically significant genes depicted in red.

B- Differentially expressed genes in SSc fibroblasts vs Control fibroblasts. Statistically significant genes depicted in red.

C- Overlap of Differentially expressed genes in SSc myofibroblasts (vs SSc nonmyogenic fibroblasts), activated myofibroblasts (vs other fibroblast subtypes), and SSc fibroblasts (versus control fibroblasts)

D- Shared differentially expressed genes that are upregulated in SSc myofibroblasts, activated myofibroblasts, and SSc fibroblasts

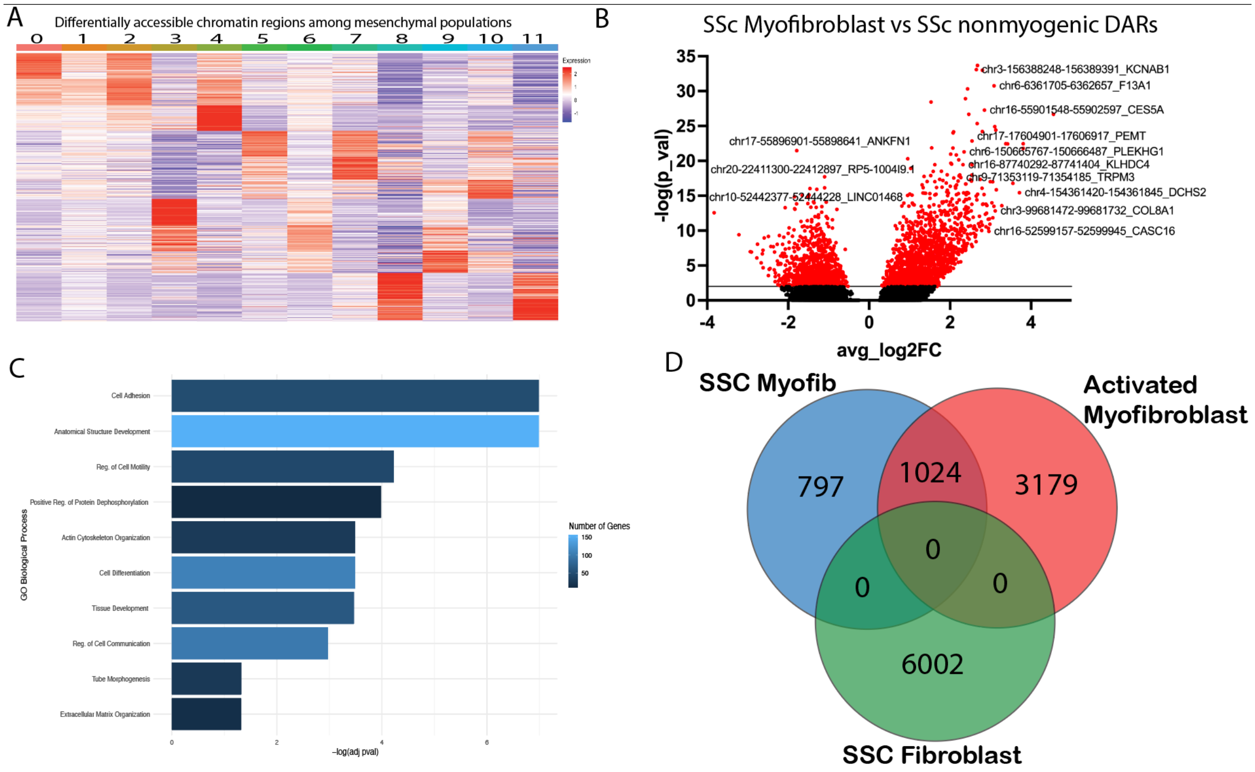


**Supplemental Figure 8**

A- Differentially accessible chromatin regions among mesenchymal cell populations.

B- SSc Myofibroblasts vs SSc Nonmyogenic fibroblasts DARs with labeling of select DARs and the nearest annotated gene.

C- Pathway enrichment analysis of chromatin regions more accessible in SSc Myofibroblasts vs SSc Nonmyogenic fibroblasts.

D- Overlap of differentially accessible regions in SSc myofibroblasts (vs SSc nonmyogenic fibroblasts), activated myofibroblasts (vs other fibroblast subtypes), and SSc fibroblasts (versus control fibroblasts)

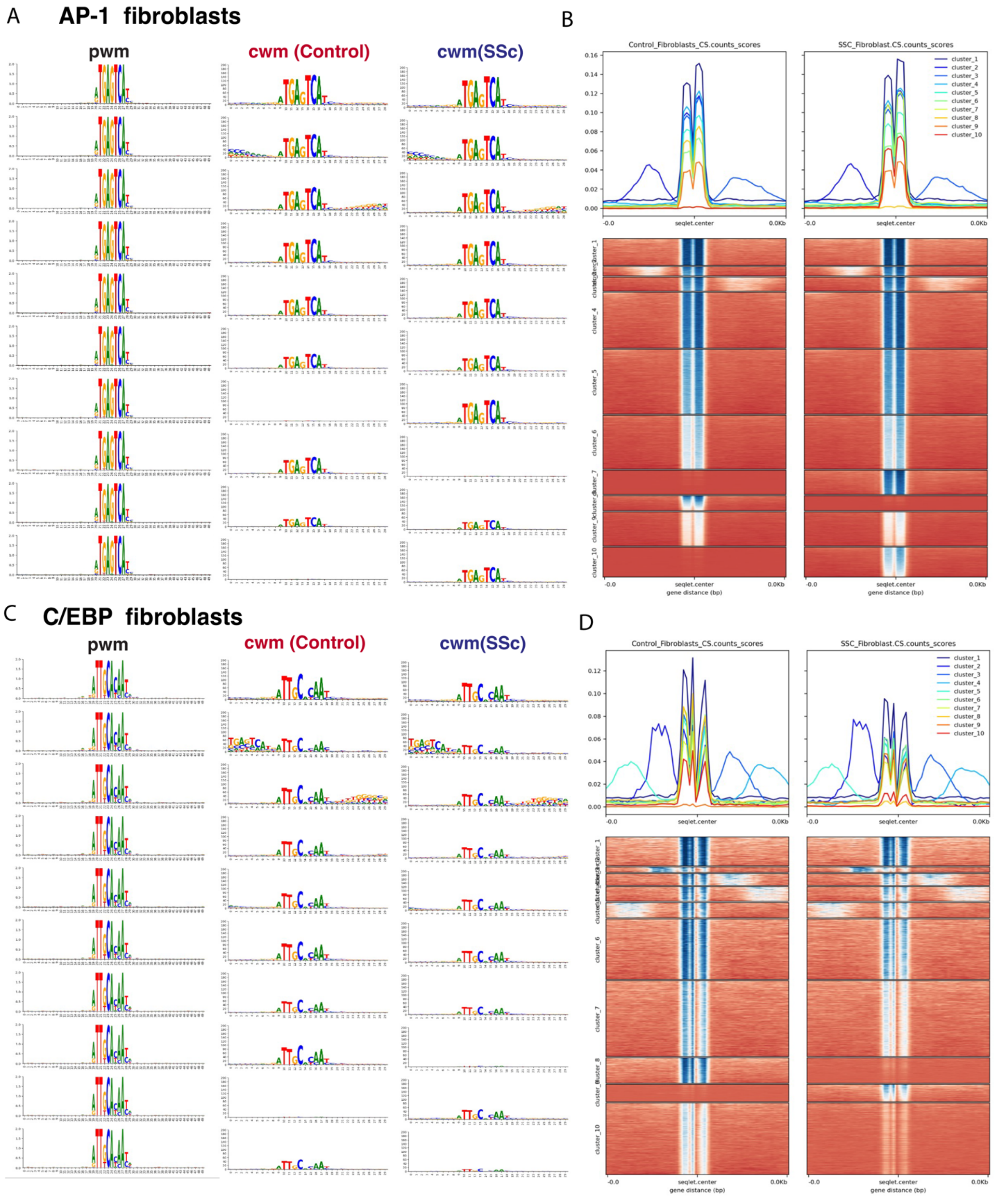


**<u>Supplemental Figure 9</u>**

A- Position weighted matrix queried and contribution weighted matrices for SSc and control fibroblasts by cluster for AP-1 seqlets.

B- Contribution score dot products (each cluster depicted in different color) and heatmap of AP-1 contribution score dot product for each seqlet/chromatin region divided by SSc and control fibroblasts into 10 clusters. Darker blue indicates higher contribution score.

C- Position weighted matrix queried and contribution weighted matrices for SSc and control fibroblasts by cluster for C/EBP seqlets.

D- Contribution score dot products (each cluster depicted in different color) and heatmap of C/EBP contribution score dot product for each seqlet/chromatin region divided by SSc and control fibroblasts into 10 clusters. Darker blue indicates higher contribution score.

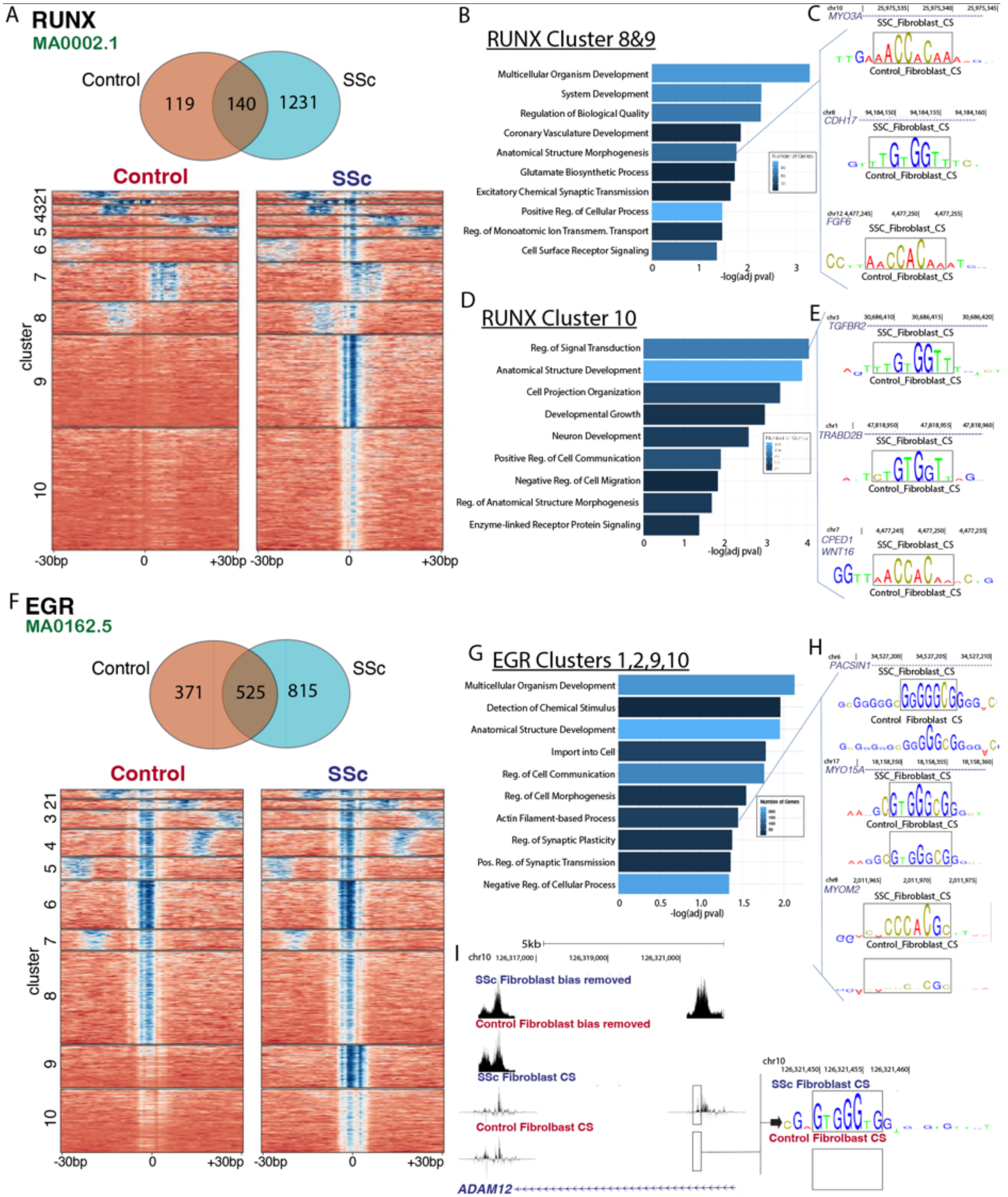


**Supplemental Figure 10**

A- Number of RUNX seqlets unique to and shared by SSc and control fibroblasts. Heatmap of RUNX contribution score dot product for each seqlet/chromatin region divided by SSc and control fibroblasts into 10 clusters. Darker blue indicates higher contribution score.

B- Pathways enriched by gene ontology for the RUNX clusters 8 and 9 chromatin regions.

C- Contribution score track depictions for SSc and control fibroblasts of 3 chromatin enriched in the anatomical structure morphogenesis pathway, from RUNX clusters 8 and 9.

D- Pathways enriched by gene ontology for the RUNX cluster 10 chromatin regions.

E- Contribution score track depictions for SSc and control fibroblasts of 3 chromatin regions (annotated to *TGFBR2, TRABD2B,* and *CPED1/WNT16*) enriched in the regulation of signal transduction pathway, from RUNX cluster 10.

F- Number of EGR seqlets (as detected for the MA0162.5 EGR1 matrix profile) unique to and shared by SSc and control fibroblasts. Heatmap of EGR contribution score dot product for each seqlet/chromatin region divided by SSc and control fibroblasts into 10 clusters. Darker blue indicates higher contribution score.
G- Pathways enriched by gene ontology for the EGR clusters 1, 2, 9, and 10 chromatin regions.
H- Contribution score track depictions for SSc and control fibroblasts of 3 chromatin regions (annotated to *PACSIN1, MYO15A,* and *MYOM2*) enriched in the regulation of actin-filament based process pathway, from EGR clusters 1, 2, 9, and 10.
I- Pseudobulk bias removed signal and contribution score tracks for *ADAM12* region with EGR footprinting in SSc fibroblasts.

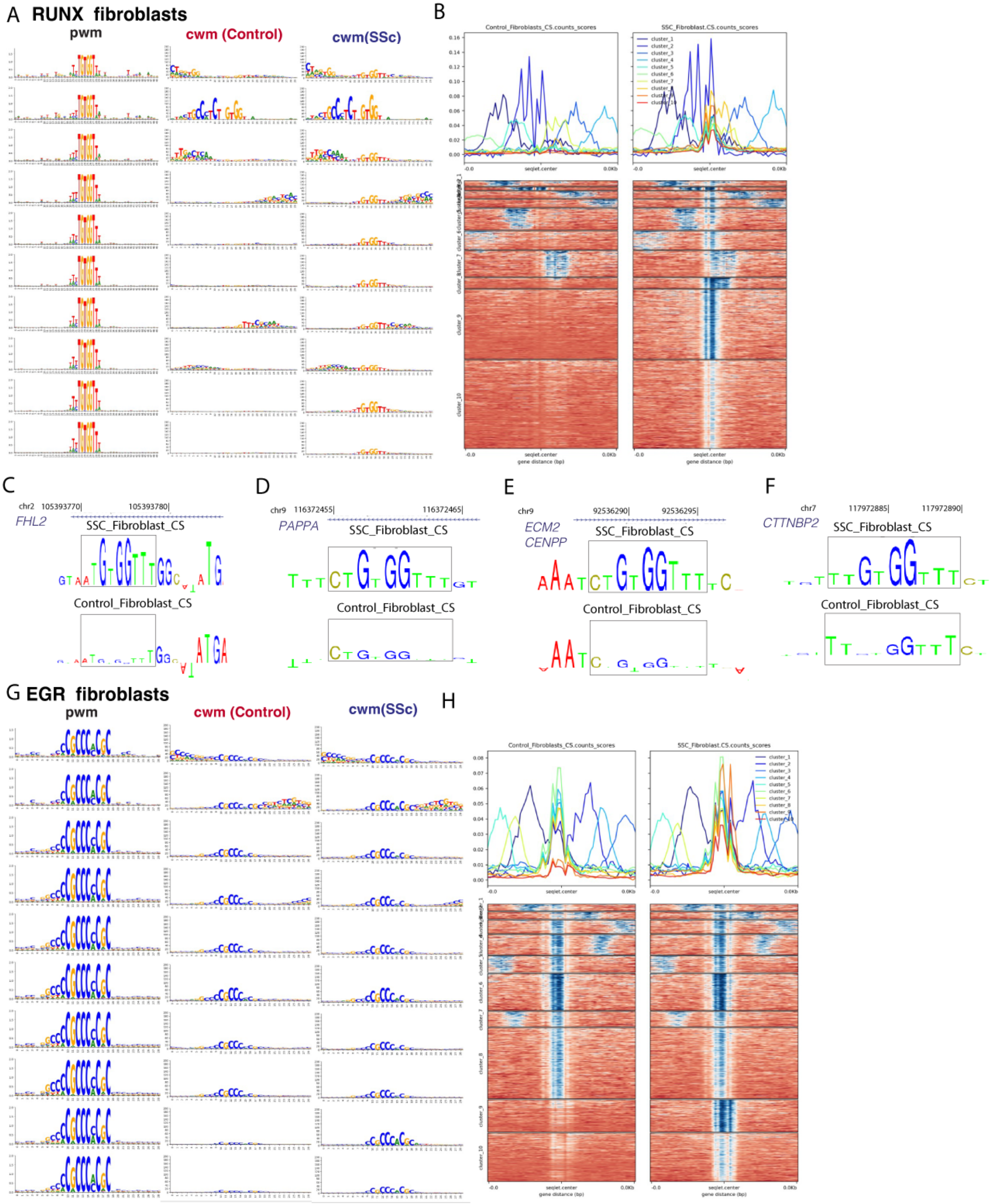


**Supplemental Figure 11**

A- Position weighted matrix queried and contribution weighted matrices for SSc and control fibroblasts by cluster for RUNX seqlets.

B- Contribution score dot products (each cluster depicted in different color) and heatmap of RUNX contribution score dot product for each seqlet/chromatin region divided by SSc and control fibroblasts into 10 clusters. Darker blue indicates higher contribution score.

C- Contribution score track depiction for SSc and control fibroblasts of region annotated to *FHL2* with enriched RUNX action in SSC fibroblasts.

D- Contribution score track depiction for SSc and control fibroblasts of region annotated to *PAPPA* with enriched RUNX action in SSC fibroblasts.
E- Contribution score track depiction for SSc and control fibroblasts of region annotated to *ECM2* with enriched RUNX action in SSC fibroblasts.
F- Contribution score track depiction for SSc and control fibroblasts of region annotated to *CTTNBP2* with enriched RUNX action in SSC fibroblasts.
C- Position weighted matrix queried and contribution weighted matrices for SSc and control fibroblasts by cluster for EGR seqlets.
D- Contribution score dot products (each cluster depicted in different color) and heatmap of EGR contribution score dot product for each seqlet/chromatin region divided by SSc and control fibroblasts into 10 clusters. Darker blue indicates higher contribution score.

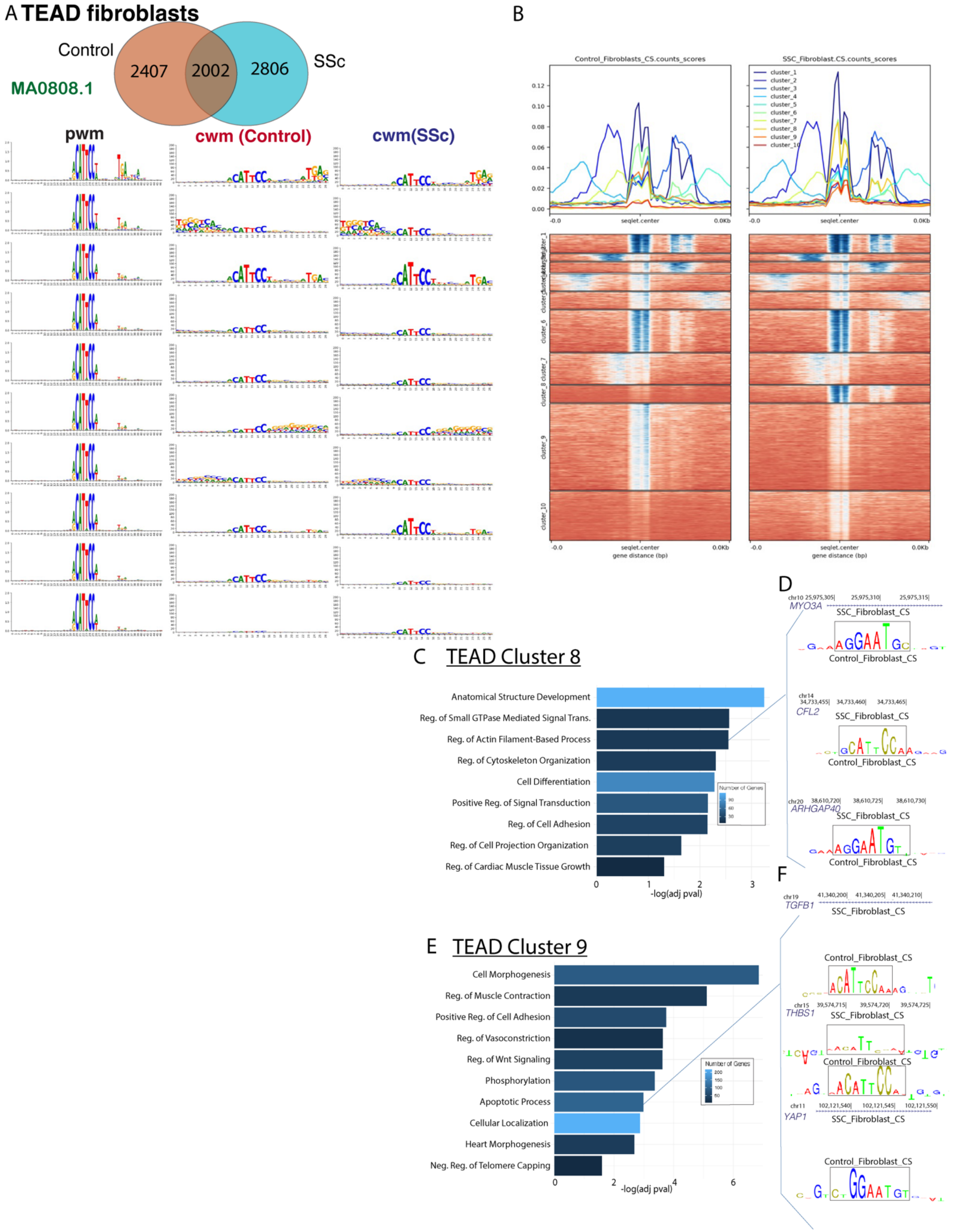


**Supplemental Figure 12**

A-Number of TEAD seqlets (as detected for the MA0808.1 TEAD3 matrix profile) unique to and shared by SSc and control fibroblasts. Position weighted matrix queried and contribution weighted matrices for SSc and control fibroblasts by cluster for TEAD seqlets.

B- Heatmap of TEAD contribution score dot product for each seqlet/chromatin region divided by SSc and control fibroblasts into 10 clusters. Darker blue indicates higher contribution score.

C- Pathways enriched by gene ontology for the TEAD cluster 8 chromatin regions (as annotated to nearest gene, limited to within 100kb).

D- Contribution score track depictions for SSc and control fibroblasts of 3 chromatin regions (annotated to *MYO3A, CFL2,* and *ARHGAP40*) enriched in the regulation of actin-filament based process pathway, from TEAD cluster 8.

E- Pathways enriched by gene ontology for the TEAD cluster 9 chromatin regions (as annotated to nearest gene, limited to within 100kb).

F- Contribution score track depictions for SSc and control fibroblasts of 3 chromatin regions (annotated to *TGFB1, THBS1,* and *YAP1*) enriched in the apoptotic process pathway, from TEAD cluster 9.

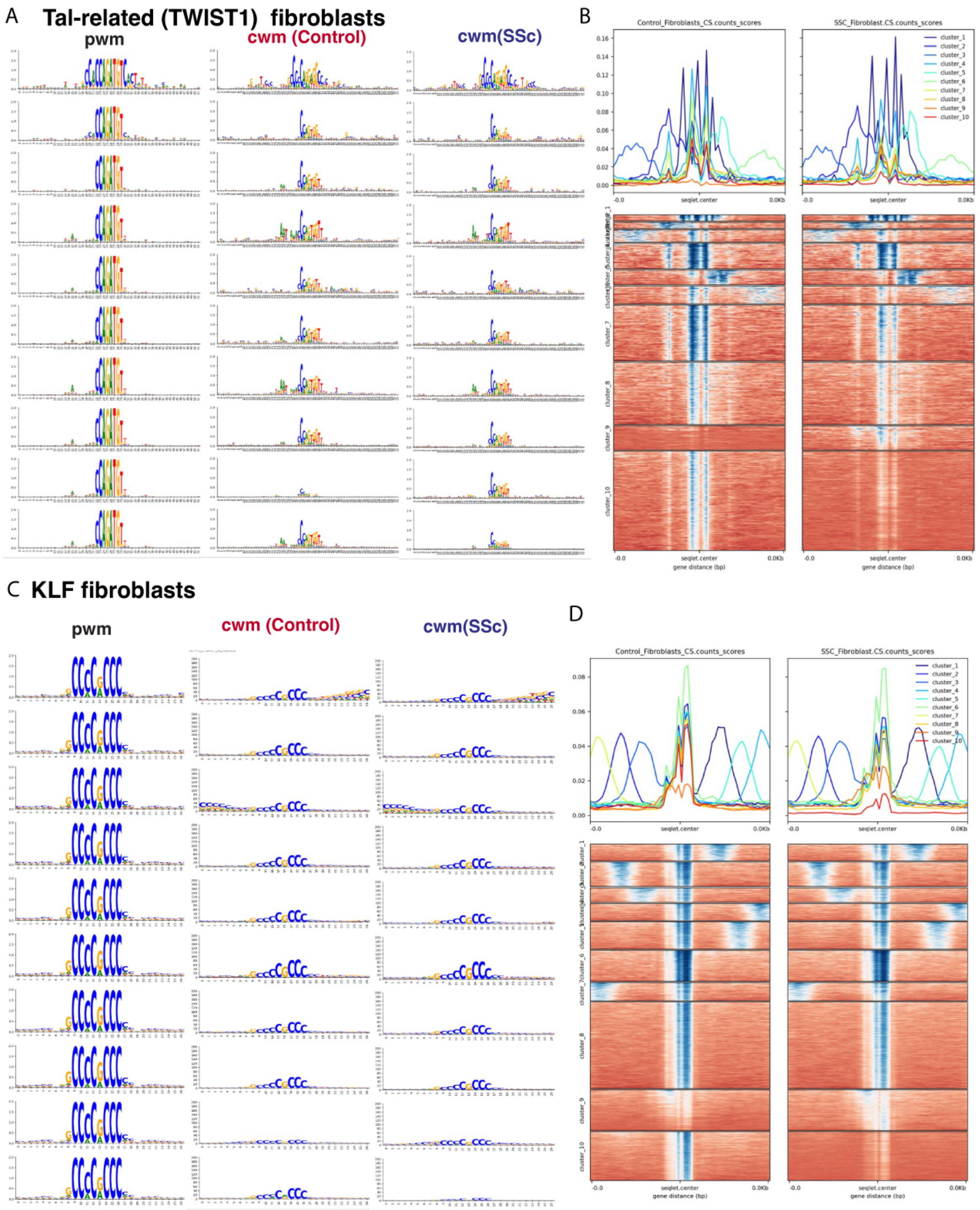


**<u>Supplemental Figure 13</u>**

A- Position weighted matrix queried and contribution weighted matrices for SSc and control fibroblasts by cluster for Tal-related seqlets.

B- Contribution score dot products (each cluster depicted in different color) and heatmap of Tal-related contribution score dot product for each seqlet/chromatin region divided by SSc and control fibroblasts into 10 clusters. Darker blue indicates higher contribution score.

C- Position weighted matrix queried and contribution weighted matrices for SSc and control fibroblasts by cluster for KLF seqlets.

B- Contribution score dot products (each cluster depicted in different color) and heatmap of KLF contribution score dot product for each seqlet/chromatin region divided by SSc and control fibroblasts into 10 clusters. Darker blue indicates higher contribution score.

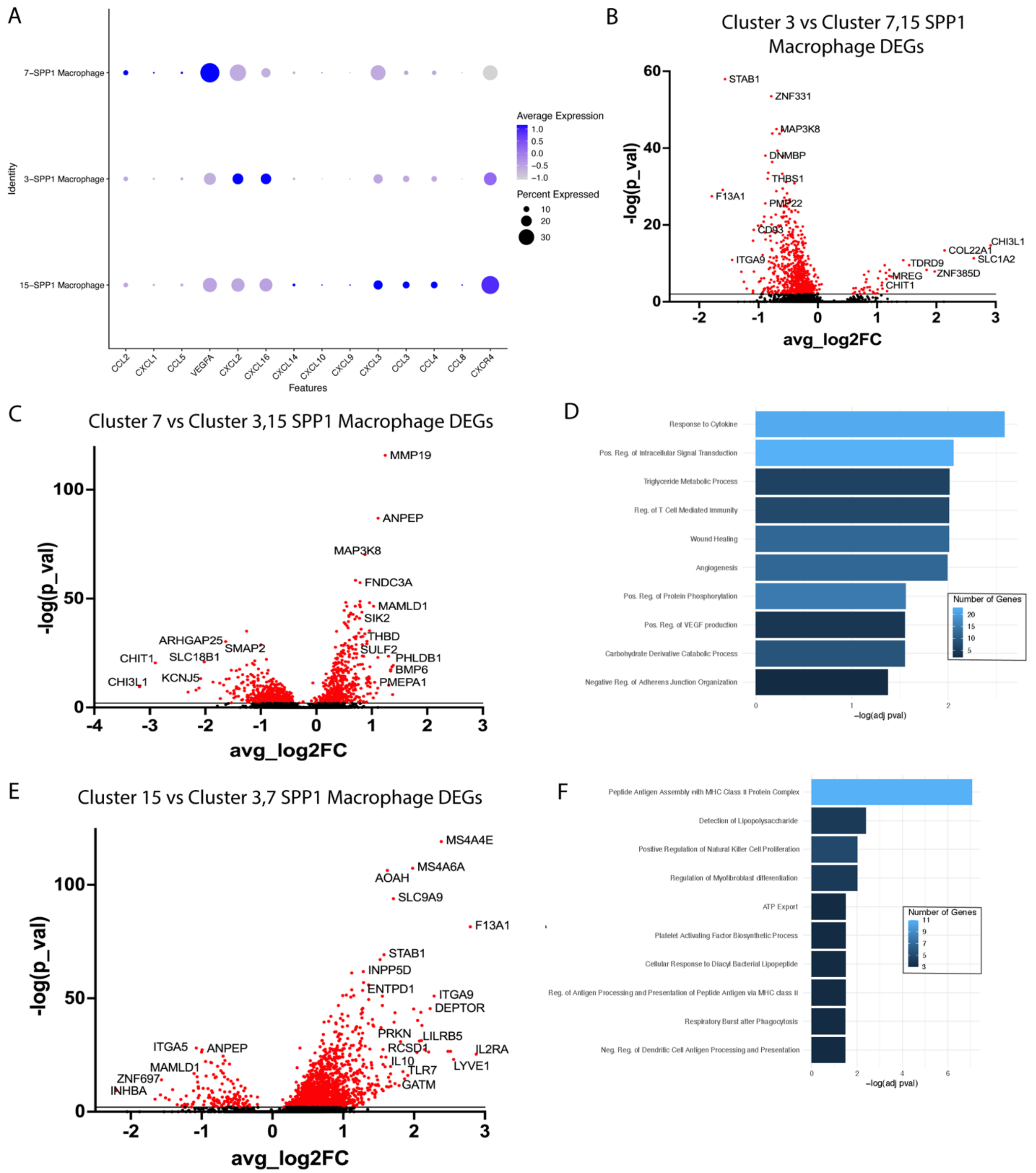


**Supplemental Figure 14**

A-Gene expression of select chemokines and growth factors in the three SPP1$^{hi}$ macrophage clusters

B-Differentially expressed genes for the Cluster 3 SPP1$^{hi}$ macrophages versus the Cluster 7 and Cluster 15 SPP1$^{hi}$ macrophages. Statistically significant genes are depicted in red.

C-Differentially expressed genes for the Cluster 7 SPP1$^{hi}$ macrophages versus the Cluster 3 and Cluster 15 SPP1$^{hi}$ macrophages. Statistically significant genes are depicted in red.

D-Pathway enrichment analysis of differentially expressed genes for the Cluster 7 SPP1$^{hi}$ macrophages versus the Cluster 3 and Cluster 15 SPP1$^{hi}$ macrophages.

E-Differentially expressed genes for the Cluster 15 SPP1$^{hi}$ macrophages versus the Cluster 3 and Cluster 7 SPP1$^{hi}$ macrophages. Statistically significant genes are depicted in red.

F-Pathway enrichment analysis of differentially expressed genes for the Cluster 15 SPP1$^{hi}$ macrophages versus the Cluster 3 and Cluster 7 SPP1$^{hi}$ macrophages.

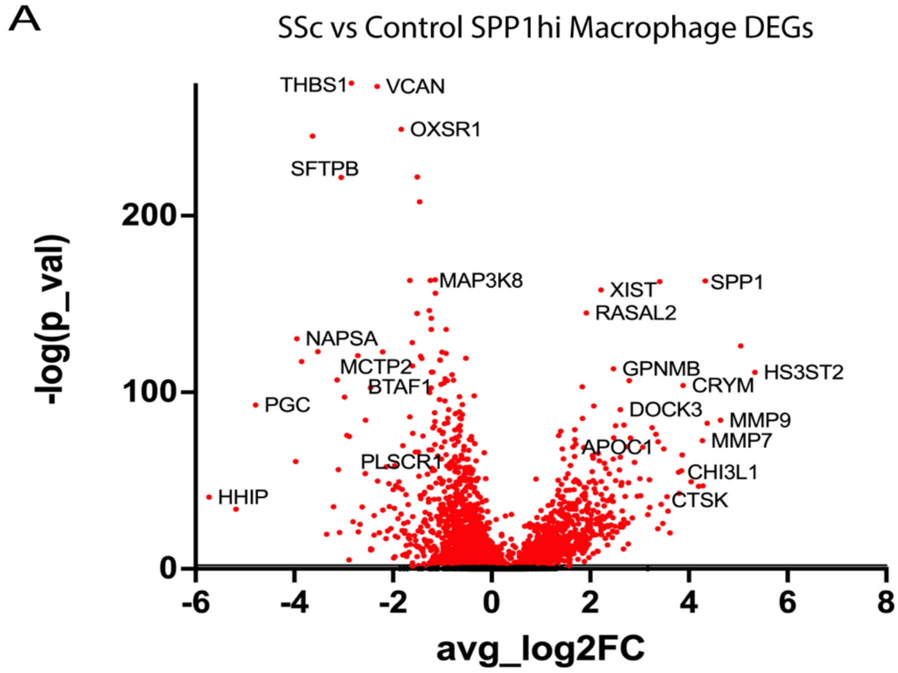


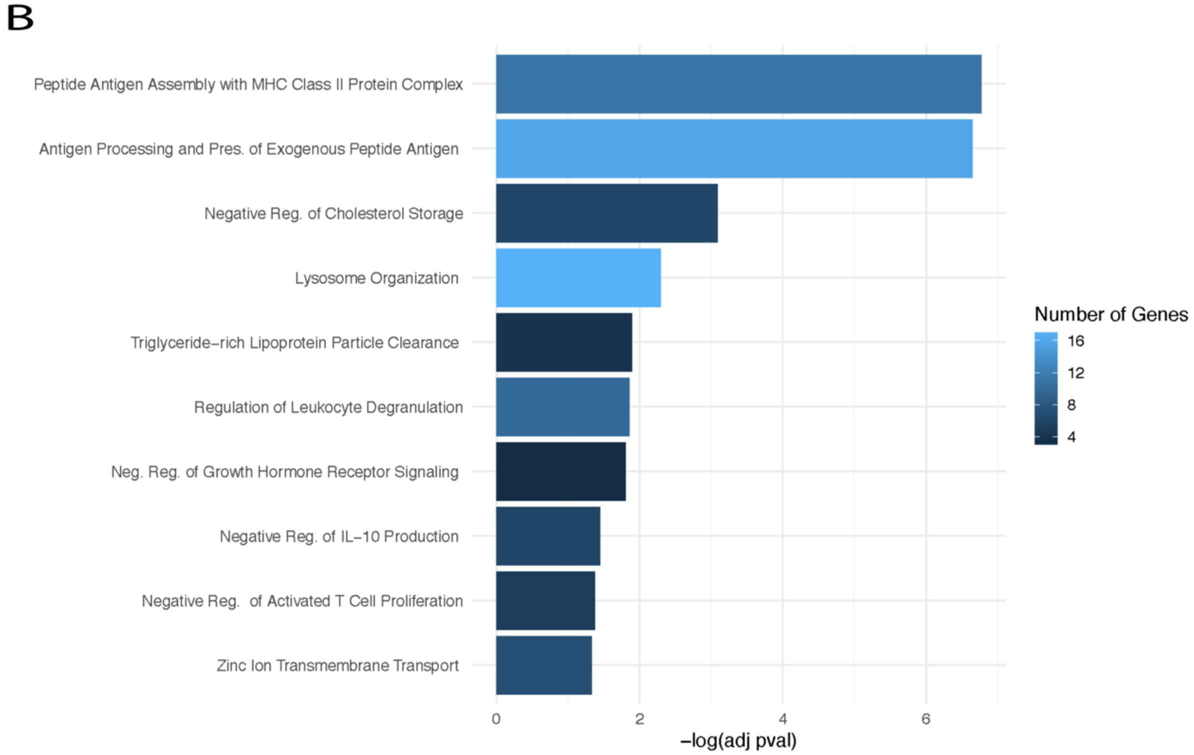


**Supplemental Figure 15**

A- Differentially expressed genes for SSc SPP1$^{hi}$ macrophages versus Control SPP1$^{hi}$ macrophages. Statistically significant genes are depicted in red.

B- Pathway enrichment analysis of differentially expressed genes for SSc SPP1$^{hi}$ macrophages versus Control SPP1$^{hi}$ macrophages.

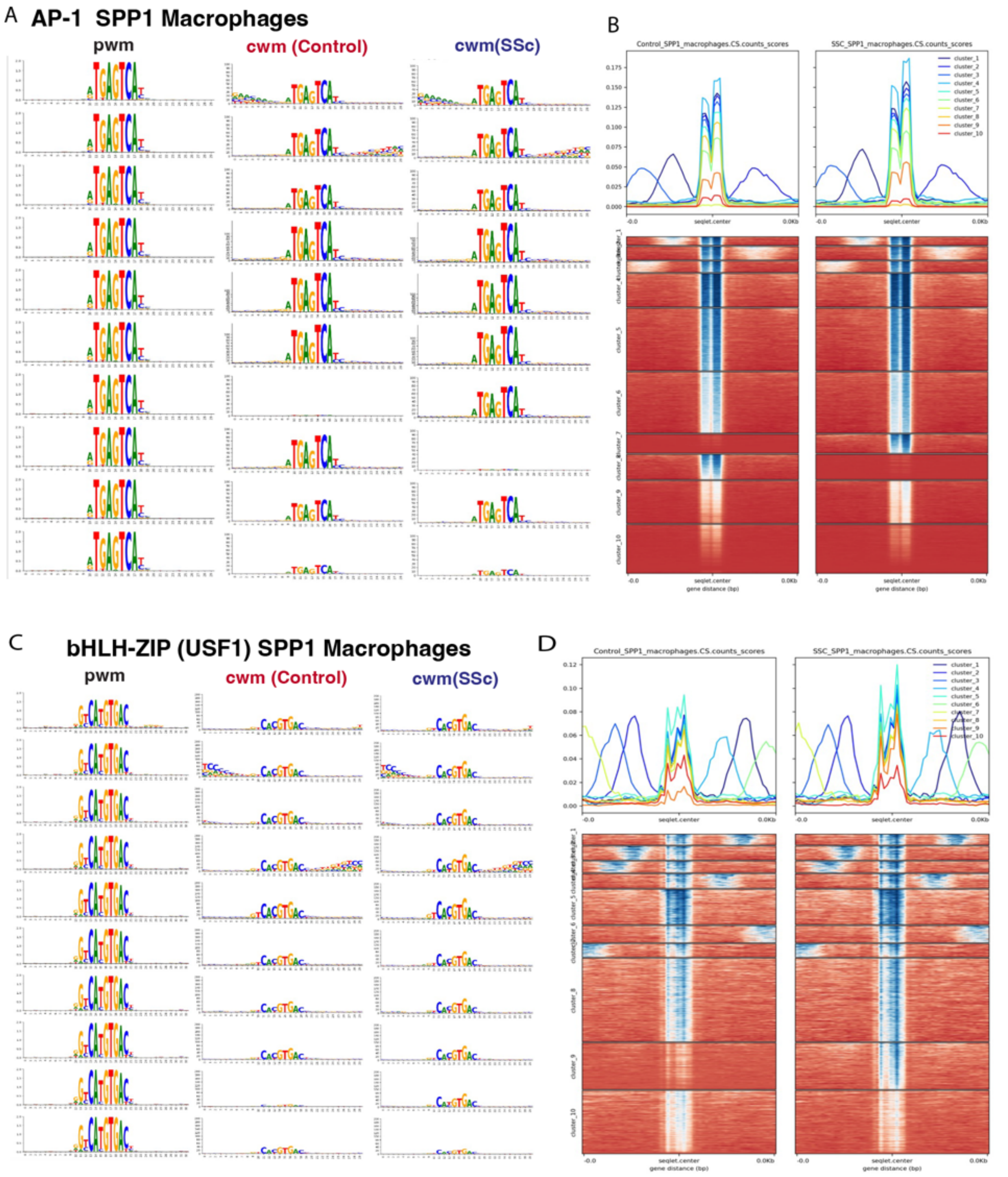


**Supplemental Figure 16**

A- Position weighted matrix queried and contribution weighted matrices for SSc and control SPP1$^{hi}$ macrophages by cluster for AP-1 seqlets.

B- Contribution score dot products (each cluster depicted in different color) and heatmap of AP-1 contribution score dot product for each seqlet/chromatin region divided by SSc and control SPP1$^{hi}$ macrophages into 10 clusters. Darker blue indicates higher contribution score.

C- Position weighted matrix queried and contribution weighted matrices for SSc and control SPP1$^{hi}$ macrophages by cluster for bHLH-ZIP seqlets.

D- Contribution score dot products (each cluster depicted in different color) and heatmap of bHLH-ZIP contribution score dot product for each seqlet/chromatin region divided by SSc and control SPP1$^{hi}$ macrophages into 10 clusters. Darker blue indicates higher contribution score.

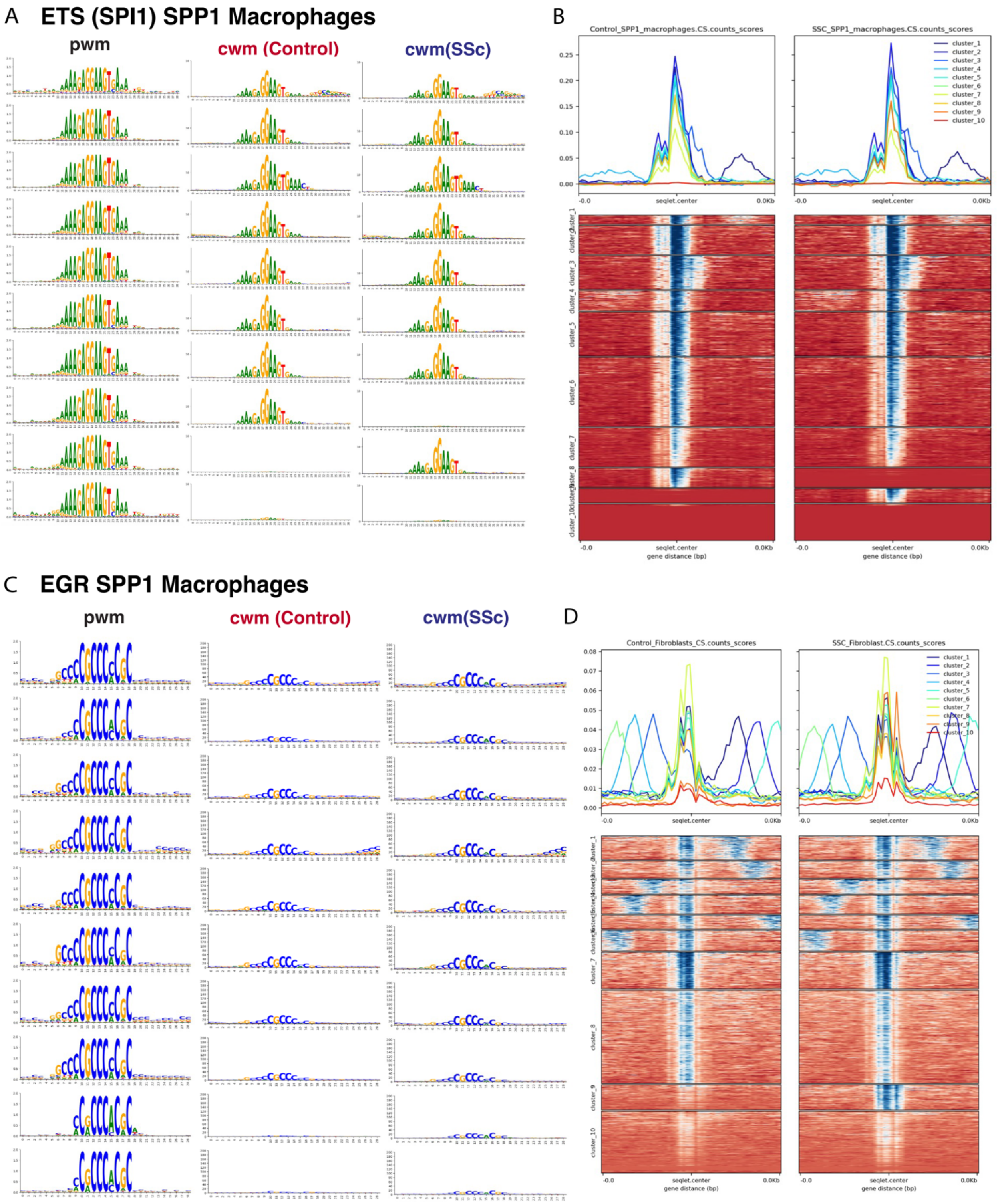


**Supplemental Figure 17**

A- Position weighted matrix queried and contribution weighted matrices for SSc and control SPP1$^{hi}$ macrophages by cluster for Ets-related seqlets.

B- Contribution score dot products (each cluster depicted in different color) and heatmap of Ets-related contribution score dot product for each seqlet/chromatin region divided by SSc and control SPP1$^{hi}$ macrophages into 10 clusters. Darker blue indicates higher contribution score.

C- Position weighted matrix queried and contribution weighted matrices for SSc and control SPP1$^{hi}$ macrophages by cluster for EGR seqlets.

D- Contribution score dot products (each cluster depicted in different color) and heatmap of EGR contribution score dot product for each seqlet/chromatin region divided by SSc and control SPP1$^{hi}$ macrophages into 10 clusters. Darker blue indicates higher contribution score.

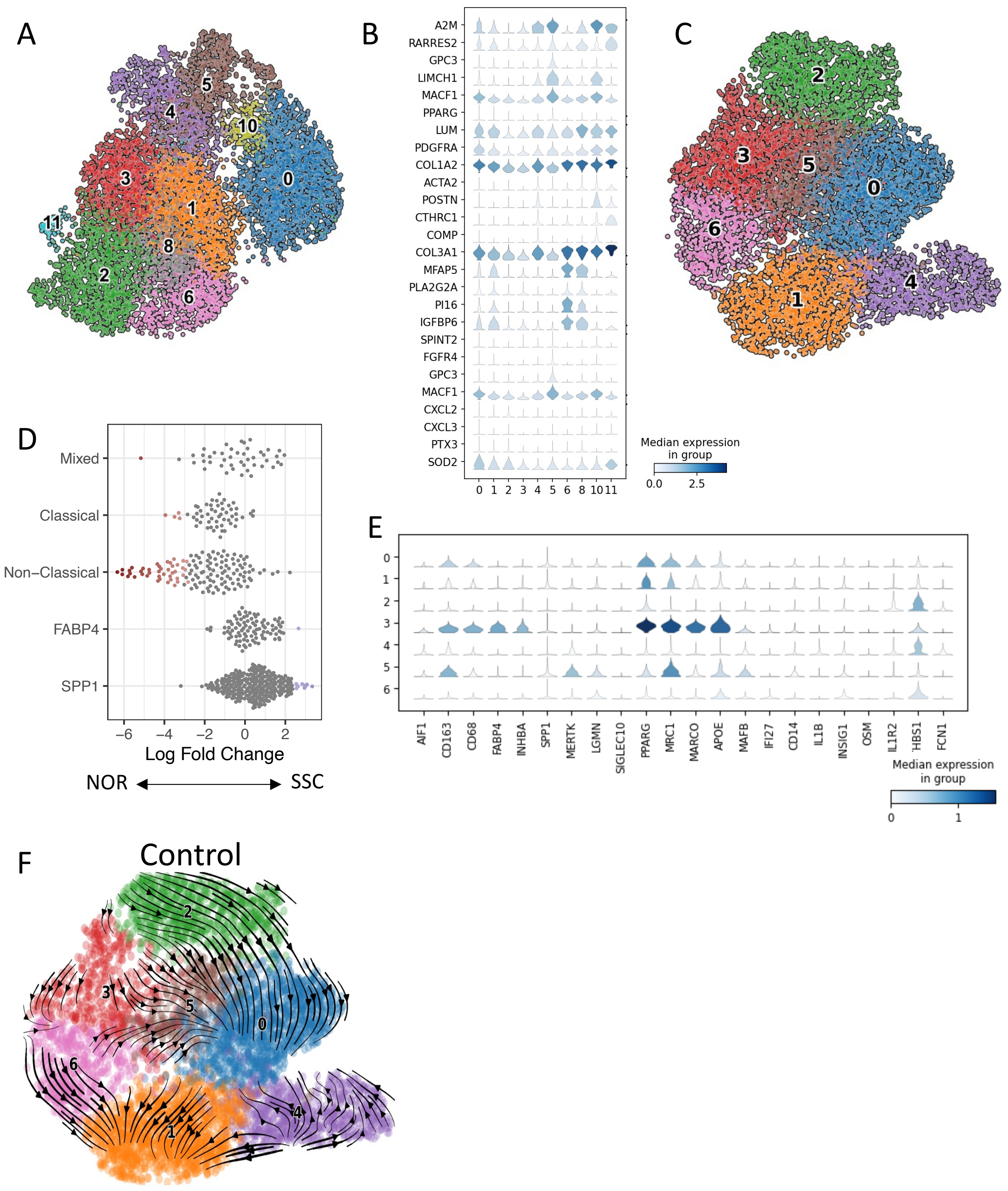


**Supplemental Figure 18**

A- Multiome samples mesenchymal clustering with smooth muscle, pericyte, and doublet clusters removed

B- Gene expression of multiome mesenchymal subclustering for select markers

C-Multiome samples myeloid clustering

D- Differential abundance testing using Milo to compare control and SSc-ILD samples by myeloid subtype.

E- Gene expression of multiome myeloid subclustering for select markers

F- RNA velocity of multiome Control myeloid cells

Supplemental Table 1

Demographics and Processing Details by Sample

| Sample | Age (year) | Sex | Race/Ethnicity | mPAP (mmHg) | Medication at time of transplant | Antibody | Chemistry | Depletions | Sample ID |
|---|---|---|---|---|---|---|---|---|---|
| SSc | 59 | F | Hispanic | 46 | ambrisentan, tadalafil | Th/To | snATAC-seq | ½ whole, ½ CD45, CD326, CD31 depletion | SC294 |
| SSc | 42 | F | Black | 27 | mycophenolate, tadalafil, macitentan, selexipag | | snATAC-seq | ½ whole, ½ CD45, CD326 depletion | SC336 |
| SSc | 61 | F | White | 42 | | centromere | snATAC-seq | No depletions | SC346 |
| SSc | 58 | F | Asian | 22 | Rituximab, tocilizumab | | Multiome | ½ whole, ½ CD45 depletion | SC369 |
| SSc | 54 | F | Black | 40 | mycophenolate | Scl70, U1RNP | Multiome | ½ whole, ½ CD163 depletion | SC381 |
| SSc | 40 | F | Hispanic | 26 | mycophenolate, rituximab, prednisone | | Multiome | ½ whole, ½ CD45, CD326, CD31 depletion | SC395 |
| SSc | 51 | M | White | 20 | mycophenolate | Scl70 | Multiome | ½ whole, ½ CD45, CD326, CD31 depletion | SC402 |
| SSc | 63 | F | White | 31 | Mycophenolate, hydroxychloroquine, ambrisentan, tadalafil | U2RNP | Multiome | STEMCELL AnnexinV live dead, 1/2 CD66 depletion, 1/2 CD66, CD326, CD45, CD31 depletion | SC430 |
| SSc | 54 | F | Black | 38 | Ambrisentan, tadalafil | | Multiome | STEMCELL AnnexinV live dead, 1/2 CD66 depletion, 1/2 CD66, CD326, CD45, CD31 depletion | SC449 |
| Control | 64 | F | | | | | snATAC-seq | No depletions | SC316 |
| Control | 38 | M | | | | | snATAC-seq | 1/2 whole, 1/2 CD45,CD326 depletion | SC324 |
| Control | 20 | F | | | | | Multiome | ½ whole, ½ CD45, CD326, CD31 depletion | SC398 |
| Control | 68 | M | | | | | Multiome | STEMCELL AnnexinV | SC421/422 |

| | | | | | | | | | |
|---|---|---|---|---|---|---|---|---|---|
| | | | | | | | | live dead ,1/2 CD66 depletion, ½ CD66, CD326, CD45, CD31 depletion | |
| Control | 20 | F | | | | | Multiome | STEMCELL AnnexinV live dead, 1/2 CD66 depletion, 1/2 CD66,CD326, CD45, CD31 depletion | SC453 |
| Control | 62 | M | | | | | Multiome | STEMCELL AnnexinV live dead,1/2 CD66 depletion, 1/2 CD66, CD326, CD45 CD31 depletion | SC466 |
| Control | 26 | M | | | | | Multiome | STEMCELL AnnexinV live dead, 1/2 CD66 depletion, 1/2 CD66, CD326, CD45 depletion | SC486 |
| Control | 25 | F | | | | | Multiome | STEMCELL AnnexinV live dead, 1/2 CD66 depletion, 1/2 CD66, CD326, CD45 depletion | SC490 |

Supplementary Files

Supplemental Figures

Supplemental Table 2- Differentially accessible regions in SSc vs control fibroblasts
Supplemental Table 3- Differentially accessible regions in SSc vs control SPP1$^{hi}$ macrophages
Supplemental Table 4-Differential gene expression, motif activity, and regulon score of transcription factors in SSc vs control fibroblasts
Supplemental Table 5- Differential gene expression, motif activity, and regulon score of transcription factors in SSc vs control SPP1$^{hi}$ macrophages
